# Retroviral adapters hijack the RNA helicase UPF1 in a CRM1/XPO1 dependent manner and reveal proviral roles of UPF1

**DOI:** 10.1101/2023.06.20.545693

**Authors:** Lea Prochasson, Makram Mghezzi-Habellah, Armelle Roisin, Martine Palma, Jean-Philippe Robin, Steve de Bossoreille, David Cluet, Maleke Mouehli, Didier Decimo, Alexandra Desrames, Thibault Chaze, Mariette Matondo, Hélène Dutartre, Maria-Isabel Thoulouze, Fabrice Lejeune, Pierre Jalinot, Stéphane Réty, Vincent Mocquet

## Abstract

The hijacking of CRM1 export is an important step of the retroviral replication cycle. Here, we investigated the consequences of this hijacking for the host. During HTLV-1 infection, we identified that this hijacking by the viral protein Rex favours the association between CRM1 and the RNA helicase UPF1, leading to a decreased affinity of UPF1 for cellular RNA and its nuclear retention. As a consequence, we found that the nonsense mediated mRNA decay (NMD), known to have an antiviral function, was inhibited. Corroborating these results, we described a similar process with Rev, the functional homolog of Rex from HIV-1. Unexpectedly, we also found that, for HTLV-1, this process is coupled with the specific loading of UPF1 onto vRNA, independently of NMD. In this latter context, UPF1 positively regulates several steps of the viral replication cycle, from the nuclear export of vRNA to the production of mature viral particles.

Graphical abstract
During retroviral replication, the nuclear export unspliced vRNA is conducted via the hijacking of the exportin CRM1 by the viral protein Rex. In parallel, the RNA helicase UPF1 is naturally exported in a CRM1 dependent manner. In the cytoplasm it drives NMD, whose substrates include vRNA. Here we demonstrated that HTLV-1 Rex dependent hijacking of CRM1 is associated with the nuclear accumulation of UPF1 and the stabilization of the interaction between CRM1 and UPF1 (1). In this complex, UPF1 shows a decreased affinity for cellular RNA associated to NMD inhibition (2). We also observed that UPF1 is selectively loaded onto vRNA and stimulates vRNA export (3). In this context, UPF1 is driven in the viral particles (without NMD cofactors) where it plays critical role in virion assembly, maturation (4) and ultimately viral infection (5). *Created in BioRender. PROCHASSON, L.* (*2025*) https://BioRender.com/urj0cvo*”*.

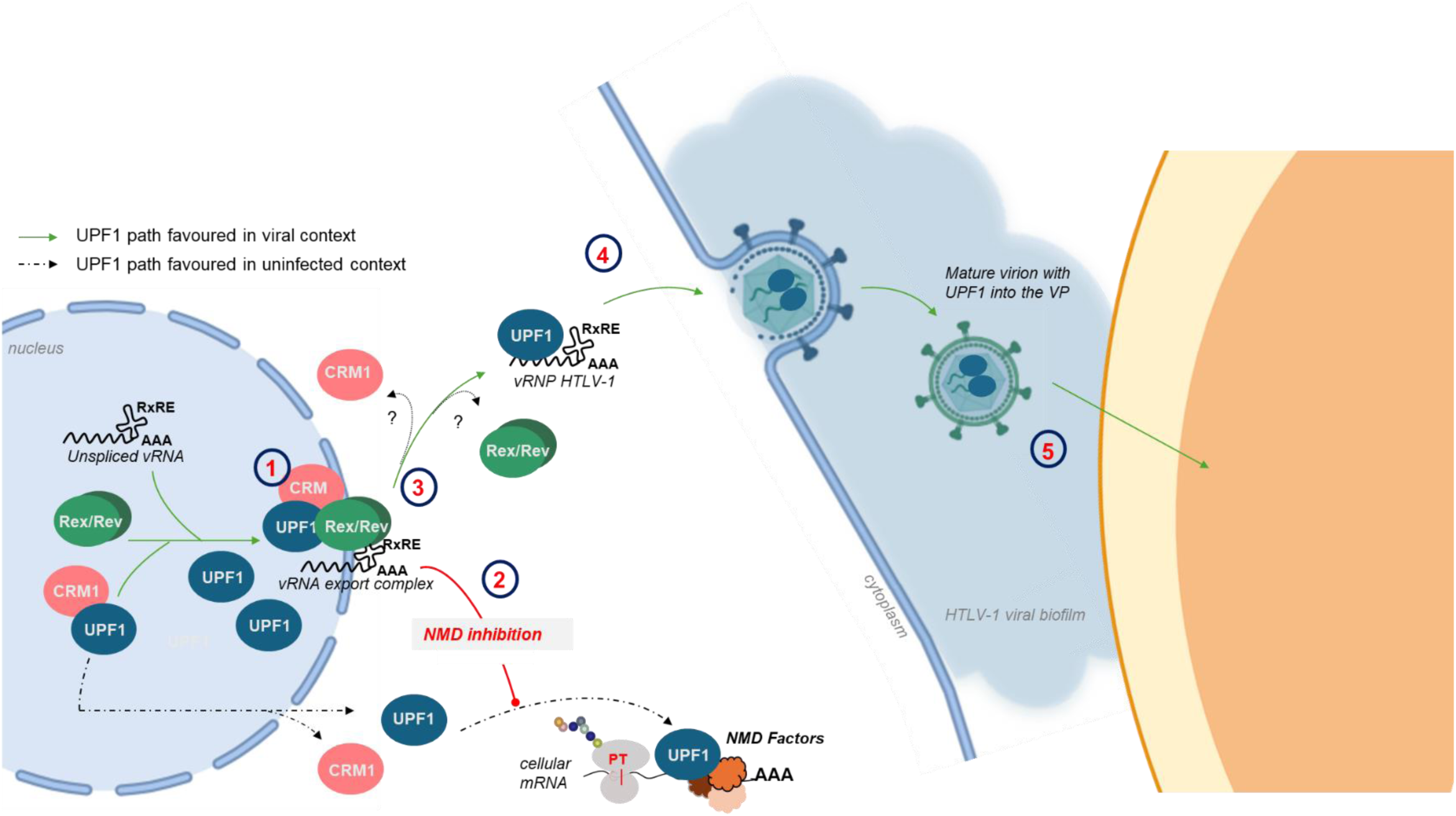

## Introduction

The maintenance of cell homeostasis requires a strict control of the repartition of macromolecules between nucleus and cytoplasm. Pathways monitoring this shuttling rely on the nature of the macromolecules to be exported or imported. Notably, whereas bulk mRNA export occurs via the NXF1/TAP1 pathway after its association with TREX complex, the exportin CRM1 (XPO1) mediates the export of proteins containing a Nuclear Export Signal (NES). These cargoes may be associated or not with a specific subset of RNA such as rRNA, snRNA and some mRNA(1, 2). NES are characterized by a specific stretch of 4 hydrophobic amino acids with variable interspaced amino acids that influences NES affinities to CRM1. The formation of the complex between CRM1 and the NES containing protein also requires the GTPase Ran to conformationally stabilize the NES binding groove of CRM1(3–6). However, the affinity of many NES substrates for CRM1 is rather low, suggesting that the formation of this trimeric transport complex is a rate-limiting step in nuclear export(7–9). At the nuclear pore, Ran GAP catalyses the conversion of RanGTP to RanGDP, inducing the dissociation of the trimeric complex CRM1/Cargo/Ran and the release of the cargo in the cytoplasm. Ultimately, the free CRM1 re-enter the nucleus for a new round of export(10). Among the plethora of functionally and structurally unrelated protein and RNP cargoes, the mislocalization of several of them due to CRM1 dysfunction supports a role of CRM1 in disease: that is the case of proteins involved in the control of DNA repair, cell cycle, transcription regulation or apoptosis such as BARD1, p21, p27, p53, FOXO3, APC and Survivin(11, 12). In addition, CRM1 overexpression and mutations were directly correlated with multiple types of cancer, notably acute myeloid leukemia and multiple myeloma(13–16). As a consequence, CRM1 was considered as a valuable target for anticancer therapies. CRM1 inhibitors where shown to sensitize cancer cells to apoptosis and several of them are currently under clinic trials(17, 18).

An important subset of CRM1 cargoes includes proteins involved in RNA processing. Notably, the DNA/RNA helicase UPF1 and its partner UPF2 have been reported to shuttle between cytoplasm and nucleus via CRM1: interactions were identified by mass spectrometry and treatments with the CRM1 inhibitor leptomycin B, induced their nuclear retention(19–21). However, the impact of a shuttling perturbation on their cellular functions have never been evaluated. UPF1 works at the crossroad of multiple processes for DNA and RNA maintenance and is directly involved in the nonsense mediated mRNA decay (NMD)(22). NMD targets and degrades mRNA harbouring premature termination codons (PTC) due to the assembly of a NMD promoting complex around UPF1, between the terminating ribosome and a downstream Exon Junction Complex(23–30). In addition to PTC containing RNA, NMD was shown to target several viral RNA, exerting an intrinsic antiviral function(31–37). Particularly, the unspliced viral RNA (vRNA) from the human delta retrovirus HTLV-1 is destabilized in a UPF1 and UPF2 dependent manner(38, 39). To counter this threat and fully express their compact genome, virus evolved different bypass strategies(40, 41). During HTLV-1 infection, the viral proteins Tax and Rex were shown to trans-inhibit NMD(38, 39). Tax directly binds UPF1 and exerts a dual inhibiting effect, repressing its ATPase activity and ultimately NMD(42). Recently, Nakano et al. showed that the N-terminal domain of Rex as well as its phosphorylation were critical for Rex/UPF1 interaction, suggesting this could lead to the incorporation of Rex in NMD complexes(43). A clear mechanism of inhibition is still to be deciphered.

Interestingly, viruses also express NES containing proteins, such as the retroviral protein Rex from HTLV-1(44). Its role in the vRNA export to the cytoplasm has been extensively studied: Rex interferes with the host splicing machinery, stabilizes vRNA over spliced isoforms and hijacks the cellular shuttling machinery to export vRNA towards translation or viral encapsidation(45–47). Notably, its Nuclear Localisation Signal (NLS) favors an importinβ−dependent nuclear translocation(48) where it accumulates in the nucleoli and binds the highly basic arginine rich motif of the vRNA called “Rex responsive element (RxRE)” (49). It also displays a dual multimerization motif encompassing the NES(50). It was proposed that Rex interacts with CRM1, multimerizes and then binds the vRNA, leading to the export of this mRNP. The cytoplasmic accumulation of vRNA resulting from the hijack of CRM1 then allows the translation of the structural precursor polyprotein GAG and the packaging of two vRNA copies within the newly synthesized viral particles(51).

While the causative role of CRM1 in disease strongly stresses the importance of a correct repartition of cargoes between nucleus and cytoplasm, understanding the impact of CRM1 hijacking by retroviral adapters on the cellular homeostasis is barely evoked in the literature. Thus, in this work, we investigated how the hijacking of CRM1 by HTLV-1 Rex affects the relationship between CRM1 and the cargo UPF1: on one hand, Rex stabilizes the UPF1/CRM1 interaction, provoking the nuclear accumulation of UPF1 and repressing its association with cellular RNA. As a consequence, this inhibits NMD, one of the cellular functions of UPF1. On the other hand, Rex drives the selective repositioning of UPF1 on the vRNA, irrespective of NMD. We found that UPF1 stimulates vRNA export and viral particles maturation, ultimately supporting the infectiveness of the infected cells. Our work thus demonstrates that Rex and CRM1 orchestrate an unexpected switch in the role of the helicase UPF1 during HTLV-1 infection.

## Methods

### Mutagenesis

To construct Rex mutant plasmids, directed mutagenesis was carried out as follow: a 20 cycles PCR was done with Phusion polymerase, 2µM of complementary primers with the point mutation and 50ng of the parental plasmid (pCMV Rex WT). PCR products were digested by *DpnI* before XL1 blue transformation. For the molecular clone mutagenesis : pCMVHTLV WT and Tax9Q plasmids were already described(52). To modify Rex, a 1kb fragment between both *BlpI* restriction sites was amplified: the 5’ forward primer harboured the nonsense mutation L23stop in the Rex sequence. It is neutral for the Tax sequence encoded on the same RNA. The DNA fragment and retroviral backbone were digested by *BlpI* (NEB). Both were ligated at 16°C overnight with T4 DNA ligase (NEB) before XL1 blue transformation. All mutants were validated by Sanger sequencing (Genewiz). See supplementary method for all the primer sequences.

### Immunoprecipitations

∼2 × 10^6^ 293T cells were transfected as described in the figures using jet prime reagent (Polyplus). 48h post transfection, cells were washed with PBS two times and pelleted in two dry pellets: 10% of the cells were put aside and directly resuspended in Laemmli buffer (loading buffer). It is referred as the INPUT fraction. The other pellet composed of the remaining 90% cells was resuspended in 400µl of lysis buffer (50 mM Tris–Cl, pH 8, 1% Triton, 10% glycerol, 0.05% SDS, 0.1 mM EDTA, 0.1 mM GTP, 200 mM KCl, protease inhibitor (Roche)) for 30 min at 4°C. The soluble fraction obtained after centrifugation at 11, 000g for 10 min was pre-cleared with sepharose beads pre-coated with tRNA and BSA-0.3% for 30 min. Supernatant was further incubated with primary antibody (5 µl) overnight at 4 °C. Protein A Sepharose® beads (Sigma) were coated with PBS + 0.3% BSA, supplemented with tRNA overnight. After re-equilibration in lysis buffer, 30 µl of beads were added to the lysate for 2h30 at 4°C before three extensive washings of 15 min in lysis buffer. The dry beads were resuspended in loading buffer for western blotting. For RNase treatment, 0.1 mg/ml RNAse A was added before the washing step. See supplementary data for uncropped images.

### Immunofluorescence

50000 cells were cultivated per well into 4 well Chamber Slide™ Labtek® II in DMEM (for HeLa or 293T cells) or RPMI medium (for Jurkat, C91PL or C8166 cells) complemented with 10% SVF and Penicillin/Streptomycin. 48h after transfection (HeLa or 293T) or after 30 minutes of sedimentation at room temperature (Jurkat, C91PL, C8166 cells), cells were fixed with 4% PFA during 20 min. After extensive washing, aldehyde groups were saturated in a 0.1M PBS Glycine solution for 30 min then washed once with PBS 1X. Cells were permeabilized with PBS-1%Triton for 5 min and then extensively washed. Saturation of non-specific sites was performed by 0.1% BSA-PBS for 30 min then extensively washed. The indicated primary antibodies were incubated for 1h30 at Room Temperature (RT) at 1/1000 (vol/vol) and further washed three times. Secondary antibodies were incubated for 50 min at 1/500 before three washes. Hoechst coloration was used to visualized nucleus (1 ng/ml final concentration). Vectashield H-1000 from Vector laboratories was used before recovering with cover slips 22*60 mm from Menzel Gläser.

### Proximity Ligation Assay (PLA)

Different combinations of plasmids were transfected in 293T or HeLa cells as indicated in the legends. PLA was carried out using the kits Duolink (Millipore Sigma) or NaveniFlex Cell MR (Navinci) and following the manufacturer indications. Dilutions of primary antibodies (vol/vol): anti-UPF1 (1/250^eme^); anti-HA (1/500^eme^); anti-FTSJ1 (1/250^eme^); anti-eIF5A (1/250^eme^) anti CRM1(1/1000eme). For UPF1/HA-Rex PLA, acquisition were performed with Spinning Disk at 60x oil objective (Live SR) microscope. Huygens Professional software was used for image deconvolution. For HA-UPF1/CRM1 PLA, image acquisitions were performed with the confocal microscope LSM800 at 20x objective with the software Zen Pro (Zeiss). ImageJ software was used to image treatment and the count was based on 8 pictures per condition. Experiments were repeated twice.

### Image acquisition, treatment and counting

Image acquisitions were performed with the confocal microscope LSM800 and a 20x or a 63x oil objective. ImageJ software (Fuji Version 2.3.0) was used to apply ‘gaussian blur’ filter (3.00 rad) to all images. Then colocalization pixel was performed on 8-bit gray scale images with the function “image calculator” and “AND” command between channels of interest. Gray scale was used to identify common pixels. To determine the % of cells with a nuclear accumulation of UPF1, we determine a threshold on the number of common pixels between UPF1 and nucleus to class cells. For the nucleus/cytoplasmic ratio quantification of UPF1 signal, the detection of cytoplasmic and nucleus regions of interest (ROIs) was determined on the signal strength from UPF1+CRM1 channels and Hoechst+CRM1 channels respectively, then manually verified and/or adjusted. To avoid quantification bias from disparate cell thickness and proximity of different cells, cytoplasmic ROI was restricted to a ring of 20 pixels width around the nuclear ROI. Then, nuclear/cytoplasmic ratio was extracted from fluorescence intensity mean in each ROI These segmentation and quantification processes have been done using Fiji macros available here: https://gitbio.ens-lyon.fr/sdebosso/mocquet/. n = 54 for control, n = 36 for Rex, n = 37 for Rev, n = 67 for C91PL and n = 66 for C8166 conditions. P values were calculated by performing first Fischer test then a Student’s t-test (paired, two-tailed) ns: P > 0.05; *P < 0.05; **P < 0.01. Z-stack analysis were performed with IMARIS (9.2.0) from Oxford Instrument.

### RNA immunoprecipitation (RIP)

∼6×10^6^ HeLa or 293T cells were transfected as described in the figures, harvested and resuspended in lysis buffer (50mM Tris–Cl, pH 7.5, 1% NP-40, 0.5% sodium deoxycholate, 0.05% SDS, 1mM EDTA, 150mM NaCl, protease inhibitor (Roche) and RNAsin (Promega)) as previously described(42). Extracts obtained after centrifugation at 12000g for 15 min were separated in 2: 10% is the input fraction and the 90% remaining are incubated with primary antibody overnight at 4 °C for immunoprecipiation. Protein A and G magnetic beads (Dynabeads, Life Technology; 5 µl each) were mixed and coated with PBS + 0.3% BSA, supplemented with tRNA and RNasin (Promega) overnight. After re-equilibration in lysis buffer (supplemented with tRNA), the beads were added to the lysate for 2h at 4°C before extensive washing in lysis buffer. The beads were resuspended in elution buffer (50mM Tris–HCl, pH 7, 0.1mM EDTA, 10mM DTT and 1% SDS).

Double RIP were performed as reported previously(42): ∼ 18×10^6^ HeLa cells were transfected as described in the figures, harvested and fixed with 0.05% formaldehyde for 20 min at room temperature. Then, 0.25M glycine was added for 5min before PBS washing. The cell pellet was resuspended in lysis buffer, and the lysate was sonicated (Bioruptor, diagenode). First, the HA tag was immunoprecipitated. The immunoprecipitation steps were carried out as described for simple RIP except that samples were incubated with Sepharose beads. Extensive washings were performed with 50mM Tris–Cl, pH 7.5, 1% NP-40, 0.5% sodium deoxycholate, 0.05% SDS, 1mM EDTA, 1M NaCl and 1M urea. The beads were resuspended in elution buffer (50mM Tris–Cl, pH 8, 1% Triton, 10% glycerol, 0.05% SDS, 0.1mM EDTA, 0.1mM GTP, 200mM KCl, protease inhibitor (Roche), RNAsin + HA peptide at 0.1 mg/ml) and incubated at 4°C for 1h, then for 10min at 30°C to ensure elution. Finally, the elution volume was increased to 400µl and Rex immunoprecipitation was performed. After extensive washing, the formaldehyde fixation was reversed by heating samples 45min at 70°C.

For all RIP experiments, the beads and the input fraction were treated with 1ml of RNAzol RT reagent for RNA extraction. RNA was quantified by qRT-PCR using the indicated primers. The levels of immunoprecipitated RNA are expressed as % input. The values represented in the graphs correspond to the mean of at least three biological replicates, and the error bars correspond to the SEM. P values were calculated by performing a Student’s t-test (unpaired, two-tailed) ns: P > 0.05; *:P < 0.05; **:P < 0.01.; ***:P< 0.005. See supplementary method for all the primer sequences.

### RNA decay assays and qRT-PCR/RT-PCR

RNA decay assays were performed to assess the stability of mRNA expressed from a β-globin reporter minigene that was either WT (GlobinWT) or with a PTC in the second exon (GlobinPTC). For this procedure, 0.5µg of GlobinPTC or 0.5µg of GlobinWT constructs were co-transfected as indicated with 0.5µg of renilla-expressing vector (unsensitive to NMD) in 0.7×10^6^ HeLa cells with jet prime reagent (Polyplus). Additional plasmids were co-transfected as indicated in the figures. After 48h, the cells were treated with DRB (100µg.ml^-1^) to block transcription for 0, 1, 3 or 4 h. Total mRNAs were extracted using the Macherey-Nagel RNA easy extraction kit and quantified by qRT-PCR using the QuantiTect SYBR Green qRT-PCR kit (Qiagen) and appropriate primers. The values represented in the graphs correspond to the mean of at least three biological replicates, and the error bars correspond to the SEM. Half-lives were calculated for each replicate, and P values were calculated by performing a Student’s t-test (unpaired, two-tailed) ns: P > 0.05; *P < 0.05; **P < 0.01.

### Preparation of viral samples for proteomic studies

CEM T cells (as a non-infected control) and HTLV-1-infected T cell line C91PL were obtained from the NIH AIDS Research and Reference Reagent Program. Each cell preparations were obtained from cultures (5×10^5^cells.ml^-1^) grown for 3 days. These preparations were performed 3 times independently.

Free particles samples preparation: Supernatants (30ml) were centrifugated (5 min at 800g) and filtered through a 0.2μm diameter pore filter (Millipore, MA) to remove cell debris and the viral biofilms potentially detached from cultured cells. Individual cell-free virions were then purified by ultracentrifugation through a 20% sucrose (wt/vol in PBS) cushion at 83, 000 g in an SW41 rotor (Beckman) for 2 h min at 4°C. Viral pellets were resuspended in serum-free RPMI 1640 medium to obtain a 500-fold concentrated suspension of individual cell-free viral particles.

Extracellular fraction samples (ECF) preparation: preparations ECF was carried out as described previously(53), with minor modifications. Briefly, C91PL and CEM cells were washed 2 times in serum-free RPMI 1640 medium before being concentrated 3 times in serum-free RPMI 1640 medium (10 ml). Cell suspensions were then gently vortexed with a gentle MACS™ Dissociator (Miltenyi Biotech). The cells were pelleted by centrifugation and supernatants containing the ECF were collected. ECF were purified by ultracentrifugation through a 20% sucrose (wt/vol in PBS) cushion at 38, 500g in an SW41 rotor (Beckman) for 1 h min at 4°C. ECF pellets were then resuspended in RPMI 1640 serum-free medium.

Three independent replicates were performed for each sample type. Prior to processing for proteomic studies, infectivity of samples was inactivated by heating at 85°C for 30 minutes in the presence of 0.5% SDS and 10mM Hepes.

### Mass spectrometry analysis

Samples were vortexed in 1% IgePal (Merck) and digested ON with PGNase F (Merck) before solubilization in 8M urea and 50 mM Tris, pH 8.0. Proteins were reduced with 5 mM TCEP for 30 minutes at room temperature, followed by alkylation with 20 mM IAM for 30 minutes in the dark. The urea concentration was reduced below 2M with 50 mM ammonium bicarbonate. Digestion was performed first with Endoprotease LysC (Promega) and then with trypsin (ratio of 1:50) at 37°C for 3 hours and overnight, respectively. The digestion was stopped with 1% formic acid (FA) before desalting using a C18 SepPak column (Waters). The eluted peptides were dried and resuspended in 0.1% FA before injection.

LC-MS/MS analysis of the digested peptides was conducted using an Orbitrap Q Exactive Plus mass spectrometer coupled to an EASY-nLC 1200 (Thermo Fisher Scientific). A *home-made* column (C18 50 cm capillary column picotip silica emitter tip (75 μm diameter) filled with 1.9 μm Reprosil-Pur Basic C18-HD resin, Dr. Maisch GmbH, Ammerbuch-Entringen, Germany) was used for peptide separation. Peptides were loaded in 0.1% FA at 900 bars and separated at 250 nl/min using a gradient of ACN, 0.1% FA, with a step-wise increase from 3% to 7% over 8 minutes, 7% to 23% over 95 minutes, and 23% to 45% over 45 minutes (total chromatographic run time of 170 minutes, including a high ACN level step and column regeneration).

Mass spectra were acquired in data-dependent acquisition mode using the XCalibur 2.2 software with automatic switching between MS and MS/MS scans using a top 12 method. MS spectra were acquired at a resolution of 35, 000 at m/z 400 with a target value of 3 × 10^6^ ions. The scan range was limited from 300 to 1700 m/z. Peptide fragmentation was performed using higher-energy collision dissociation with an energy setting of 27 NCE. The intensity threshold for ion selection was set at 1 × 10^6^ ions with charge exclusion for z = 1 and z > 7. The MS/MS spectra were acquired at a resolution of 17, 500 at m/z 400, and the isolation window was set at 1.6 Th. Dynamic exclusion was employed within 45 seconds.

Data analysis was conducted using MaxQuant (version 2.1.4.0) with the Andromeda search engine (54,55) against a reference proteome of Human 9606 (22811 entries, downloaded from UniProt on May 10th, 2023) and HTLV downloaded from uniprot). The following search parameters were applied: carbamidomethylation of cysteines was set as a fixed modification, oxidation of methionine and protein terminal acetylation were set as variable modifications. The mass tolerances in MS and MS/MS were set to 5 ppm and 20 ppm respectively. Maximum peptide charge was set to 7 and 5 amino acids were required as minimum peptide length. A false discovery rate of 1% was set up for both protein and peptide levels. The iBAQ intensity was used to estimate the protein abundance within a sample. The match between runs features was allowed. For the relative quantification between conditions (C91PL vs CEM) only proteins present in minimum two replicates of each condition were considered for further analysis. Then, the IBAQ value of each protein was normalized by CD81, which is considered as an exosome marker. Fold Change (IBAQ C91PL/IBAQ CEM) was determined and statistical analysis. (unilateral T.test) was performed to obtain p-value. Plot were generated using ggplot2 packages (RStudio Version 2023.03.0+386 (2023.03.0+386)).

### Viral transmission assay

Viral transmission assay were performed as previously described (53, 56) with the following modifications : 0.2×10^6^ C91PL were transduced at J0 for 24h with lentivirus expressing shRNA against UPF1 or a Control (Dharmacon) at MOI =1 in 1ml of RPMI. J4: shRNA expression was induced with doxycycline treatment (0.5µg/ml). To ensure a maximum of viral biofilm renewal in the absence of UPF1, a second dose of doxycycline treatment was applied at J7. At J10, C91PL were counted: 0.05×10^6^ cells were co-cultured with 0.15×10^6^ Jurkat cells engineered to stably express luciferase under the control of a HTLV-1 LTR promoter (selection with 450ng/ml hygromycin). In parallel, remaining C91PL were analysed by FACS to monitor extracellular levels of ENV, indicative of surface-bound viral particles levels (viral biofilm). After 24h of co-culture (J11), cells were treated for Luciferase assay, following the manufacturer protocol (Promega). Luciferase values were normalized by the corresponding ENV levels, previously quantified. This normalization was possible since we identified that ENV levels were weakly affected by UPF1 knock down. The graphs represent the mean of at least 3 independent viral transmissions with SEM, and P values were calculated by performing a Student’s t-test (unpaired, two-tailed) ns: P > 0.05; ***P < 0.005.

## Results

### 1. Rex interferes with CRM1-dependent export of UPF1

In order to investigate the consequences of CRM1 hijacking by Rex, the cellular distribution of a mCherry, encompassed with a NLS and a NES, was examined by confocal microscopy. The chimeric mCherry is considered as a model of shuttling protein, with a strong NES and an intermediate NLS (57). While the mCherry was distributed among nucleus and cytoplasm with a more frequent cytoplasmic accumulation, the overexpression of Rex led to a significant nuclear retention, as indicated by the nucleo-cytoplamsic ratio measurement (**Fig1 A and B**). To confirm this result with an endogenous protein, we looked at the RNA helicase UPF1, which nuclear export relies on CRM1 as well ((19–21) and **supplementary Fig1A and B**). Confocal microscopy experiments showed that in control condition, endogenous UPF1 is mostly cytoplasmic. Strikingly, when Rex is expressed, UPF1 is significantly accumulated in the nucleus compared to control conditions. As shown by colocalizing pixel analysis, Rex, CRM1 and UPF1 signals overlap in the nucleus and at the periphery of the nucleus (**Fig 1C (i) and supplementary Fig2A**). The quantification of the nucleo/cytoplasmic repartition of UPF1 signal and the percentage of cells with UPF1 nuclear signal in both populations of cells indicate a nuclear retention of UPF1 upon Rex expression **(Fig 1C(ii))**. To control the specificity of Rex effect, HA-UPF1 distribution was also assessed with or without NLS-mCherry-NES expression: we couldn’t detect the nuclear retention as when Rex is expressed (**supplementary Fig2B**). Based on these observations, Rex effect is specific and not due to a competitive quenching of CRM1 at the expense of its cargoes.

**Figure 1:**
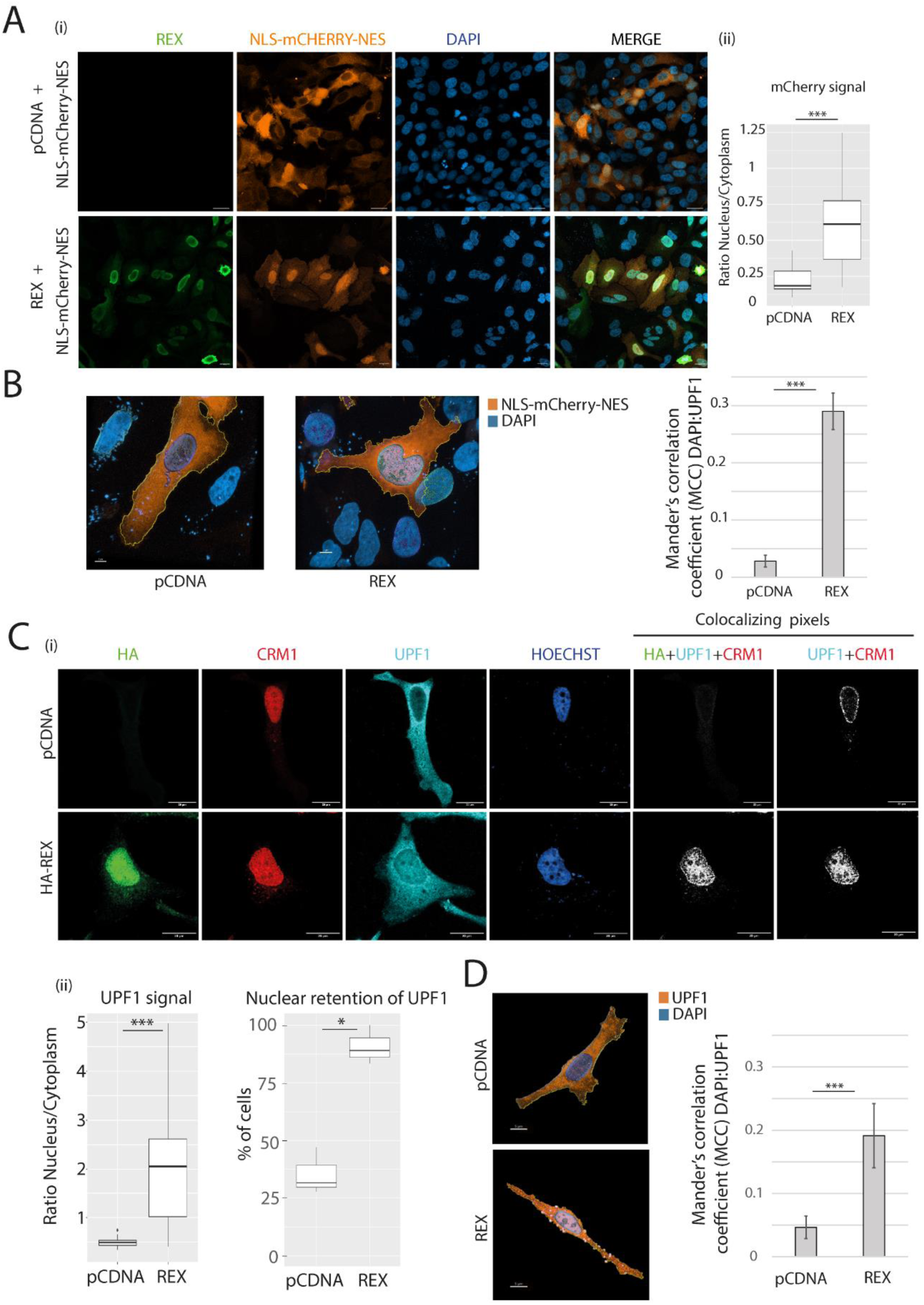
HTLV-1 Rex alters the repartition of NES containing cargos, including the RNA helicase UPF1. A) Confocal microscopy experiments were performed in HeLa cells transfected with the plasmids indicated on the left side. The factors revealed are indicated on the top (panel (i)). Objective x20. Scale bar: 20µm. Quantitative analysis of the microscopy experiments (panel (ii)): mCherry signal was quantified in the cytoplasm and in the nucleus and expressed as a ratio. Wilcoxon, pvalue ***< 0.005. B) Z-stack analysis performed with IMARIS software on images from A with mCherry and a staining of the nucleus by DAPI. The nuclear localisation of mCherry is evaluated with the Mander’s correlation coefficient (MCC). Cells were obtained from 3 independent experiments. t-test pvalue ***<0.005. Objective x63. Scale bar: 7µm. C) Confocal microscopy experiments were performed in HeLa cells transfected with the plasmids indicated on the left side (panel (i)). The factors revealed are indicated at the top. Objective x63. Scale bar: 20µm. Colocalizing pixels view is obtained as mentioned in the method section and only show the pixels simultaneously activated with the indicated channels. Quantitative analysis of the microscopy experiments (panel (ii)): on the left, UPF1 signal was quantified in the cytoplasm and in the nucleus and expressed as a ratio. Wilcoxon, pvalue *** < 0.005; on the right, the percentage of Rex expressing cells that display nuclear retention of UPF1 is indicated. t-test pvalue *<0.05. D) Z-stack analysis performed on Hela cells expressing or not Rex (without tag); UPF1 and the nucleus were stained. The nuclear localisation of UPF1 is evaluated with the Mander’s correlation coefficient (MCC). Cells were obtained from 3 independent experiments. t-test pvalue ***<0.005. Objective x63. Scale bar: 5µm.

To investigated the molecular mechanism underlying this nuclear retention of UPF1, we looked at the intracellular distribution of Rex and UPF1 with Proximity Ligation Assay (PLA), a 40nm resolution technique that detects protein-protein colocalization. In 293T cells expressing HA-Rex, PLA between Rex and endogenous UPF1 displayed numerous foci in the cytoplasm as well as in the nucleus (**Fig 2A and supplementary Fig3A**). As control, no foci were found with FTSJ1, that is not a partner of Rex, while a similar amount of foci were found with eIF5A known to interact with Rex. Those observations strongly suggest that Rex and the endogenous UPF1 can interact in the nucleus. Then, to further address this matter, we examined the effect of Rex on the formation of CRM1-UPF1 complexes, defined above to control UPF1 shuttling. Here, for a reason of antibody compatibility with the PLA kit, we used ectopic HA tagged UPF1 shown to display nuclear retention as endogenous UPF1 after Rex expression (**supplementary Fig2B)**. Strikingly, after Rex expression, PLA experiments revealed significantly more CRM1-UPF1 foci, especially in the nucleus.

**Figure 2:**
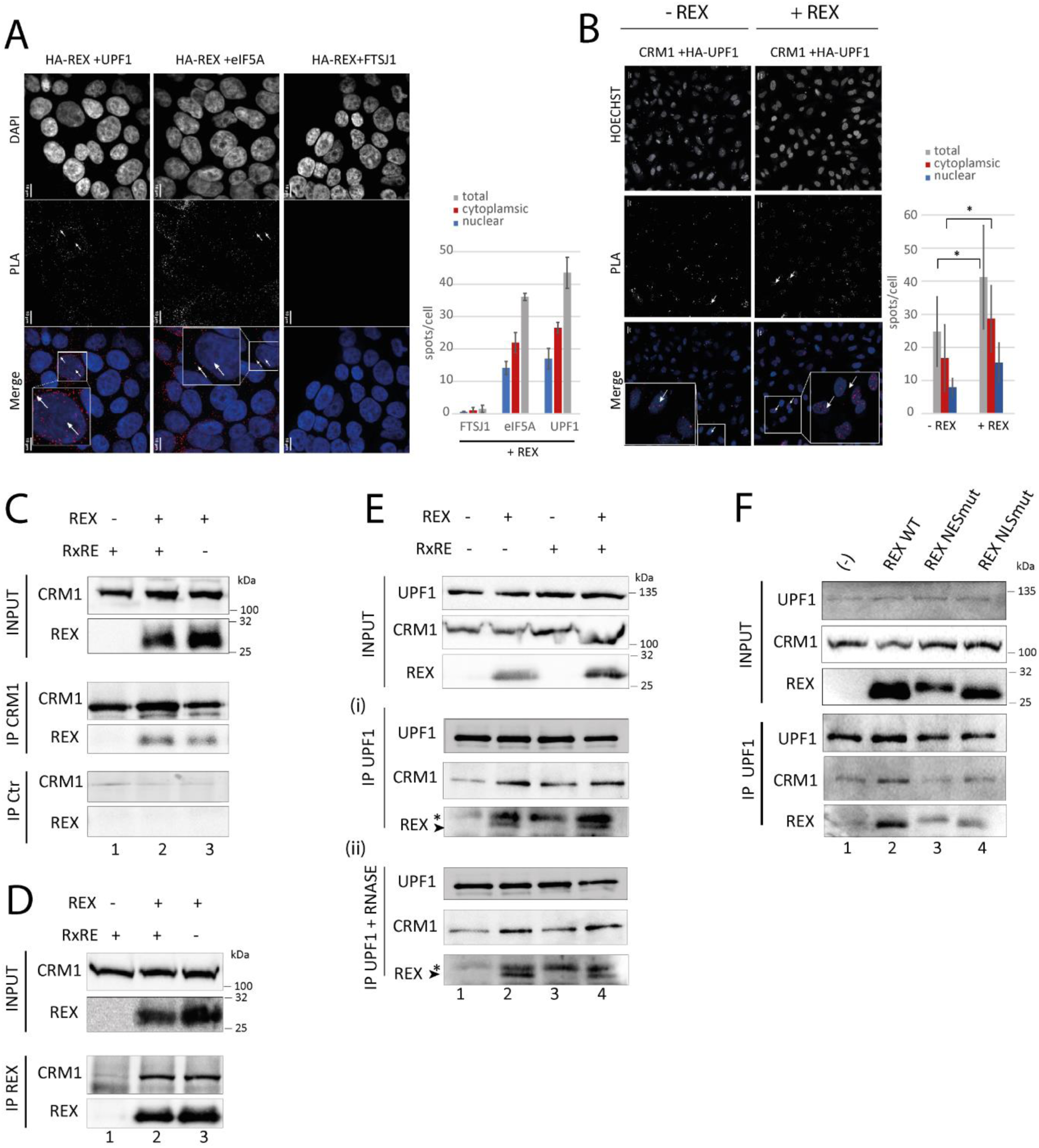
Rex interacts with CRM1 and UPF1 leading to an enhanced UPF1/CRM1 interaction and the modification of UPF1 localisation. A) Proximity Ligation Assay (PLA) carried out in 293T cells transiently transfected with a HA-REX coding plasmid. PLA combined antibodies specific of the HA tag and FTSJ1, eIF5A or UPF1 were used as indicated. FTSJ1 (ribosomal RNA methyltransferase 1) is a nucleolar protein used as a negative control (no described interaction with Rex). eIF5A, a known partner of Rex, was used as positive control. Each dot corresponds to the colocalization of the indicated protein with a resolution of 40nm. Magnification of one representative cell is framed in the merge. Two examples of dots are pinpointed by a white arrow. For each condition, the number of spots per cell and their localisation in the cytoplasmic or nuclear compartment are represented in a bar plot on the right. Objective x60. Scale bar: 10µm. B) PLA experiment carried out in HeLa cells transiently transfected with HA-UPF1 with or without Rex. PLA combined antibodies specific of the Ha Tag and endogenous CRM1. Objective x20. Scale bar: 20µm C) Co-immunoprecipitation (coIP) experiments on 293T cells extracts. Cells express the indicated combinations of Rex and RNA containing the RxRE motif. IP were performed using rabbit polyclonal antibodies against CRM1 (middle panel) or from a pre-immune serum. Proteins revealed by western blot were indicated on the side. D) Same as B with an antibody targeting Rex. E) Same as C) with an antibody targeting UPF1. CoIP were incubated for 30 min with RNAse A (0.1mg/ml) (panel (ii)) or not (panel (i)). * indicates an aspecific signal. Rex specific signal corresponds to the lower band. F) coIP experiments in 293T cells transfected with plasmids coding the indicated forms of Rex and revealed by western blot. An empty vector is transfected in lane 1. IP was performed with a rabbit polyclonal antibody against UPF1.

To confirm that Rex induced the accumulation of UPF1-CRM1 complexes, 293T cells were transfected with combinations of a plasmids encoding Rex and a RxRE containing RNA, to mimic the viral RNA export via Rex and CRM1(58). First, Rex and CRM1 co-immunoprecipitations (coIP) confirmed that Rex binds specifically to endogenous CRM1, regardless of the expression of the RxRE RNA (**Fig 2B-C)**. This supports the hypothesis that Rex can hijack CRM1 before its binding to viral RNA. Second, we revealed that endogenous UPF1 is able to coIP CRM1 and Rex. As expected, the amounts of CRM1 associated with UPF1 are increased when Rex is expressed, confirming the previously observed accumulation or stabilization of the CRM1-UPF1 complex by Rex (**Fig 2D panel (i)**). Moreover, the co-expression of the RxRE containing RNA and/or RNAse treatments did modify neither the UPF1-Rex association nor the UPF1-CRM1 enhanced interaction (**Fig 2D panel (i) and (ii)**). This indicates that the enhanced association of UPF1 and CRM1 exists independently of cellular and viral RNA. Finally, we confirmed the role of Rex in this UPF1-CRM1 association by using 2 Rex mutants (on the NES and the NLS domains). We found that the normal levels of CRM1/UPF1 association are rescued in both Rex mutants’ conditions compared to Rex WT (**Fig 2E** and quantifications in **supplementary Fig3B-D**).

Here we found that Rex expression provokes UPF1 nuclear retention as well as the accumulation or stabilisation of UPF1-CRM1 complexes, in the nucleus as well as in the cytoplasm; Altogether, it suggests that Rex alters the physiological flux of UPF1 bound to CRM1 by favouring UPF1-CRM1 interaction.

### 2. CRM1 hijacking by Rex interferes with UPF1 function during NMD

To evaluate whether the enhanced association between UPF1 and CRM1 can affect UPF1 functions, we decided to model their complex. First, while the NES of UPF1 was roughly characterized to date, we precisely delineated it, upstream the CH domain between positions 89 and 103 (**supplementary Fig4A-B**). Then, we performed 2 molecular dynamic models based on the structural representation of CRM1 interacting with a NES peptide (PDB 3GJX) and the 2 different resolved structures of UPF1: the closed form, a configuration with the CH domain positioned on the RecA2 domain (PDB 2XZL) and the open form, a more relaxed configuration with the CH domain flipped out on the RecA1 side (PDB 2WJV). After 1000ns of simulation, the UPF1 open form complexed with CRM1 was stabilized in a more compact configuration (according to the radius of gyration) and with a larger buried surface area (BSA) than the UPF1 closed form (**Fig 3A, supplementary Fig4C-D and supplementary Movies1 and 2**). The BSA, which measures the size of the interface in a protein-protein complex, is directly proportional with affinity. Moreover, it was recently suggested that in the open form of UPF1, unstructured regions are inserted in the RNA-binding channel, rationalizing that open UPF1 has a low RNA binding efficiency (59). Therefore, we suspected that when Rex is expressed, the enhanced interaction between CRM1 and UPF1 is more susceptible to involve the open conformation of UPF1, leading to a decreased affinity for RNA. To confront this hypothesis, we carried out RNA immunoprecipitation (RIP) assays by UPF1: HeLa cells were transfected with a plasmid coding a reporter mRNA (Globin PTC, a globin minigene with a premature termination codon in the second exon (60)) as well as Rex coding plasmids. UPF1 was further immunoprecipitated and the amounts of bound Globin PTC were quantified by RTqPCR. Rex expression as well as UPF1 immunoprecipitations were checked by western blot. When Rex is expressed, the level of Globin PTC associated to UPF1 is significantly reduced compared to conditions without Rex (**Fig 3B**, **supplementary Fig5A**). In addition, to correlate this effect with the previously described UPF1-CRM1 complex, we also carried out UPF1 RIP experiments in cells transfected with Rex NES mutant. As expected, while the Rex NES mutant couldn’t enhance the interaction between UPF1 and CRM1, it did not affect UPF1 binding capacity to globin PTC either (**Fig 3B**). This recovery confirms that CRM1 hijacking by Rex leads to a decreased association of UPF1 to its RNA substrate.

**Figure 3:**
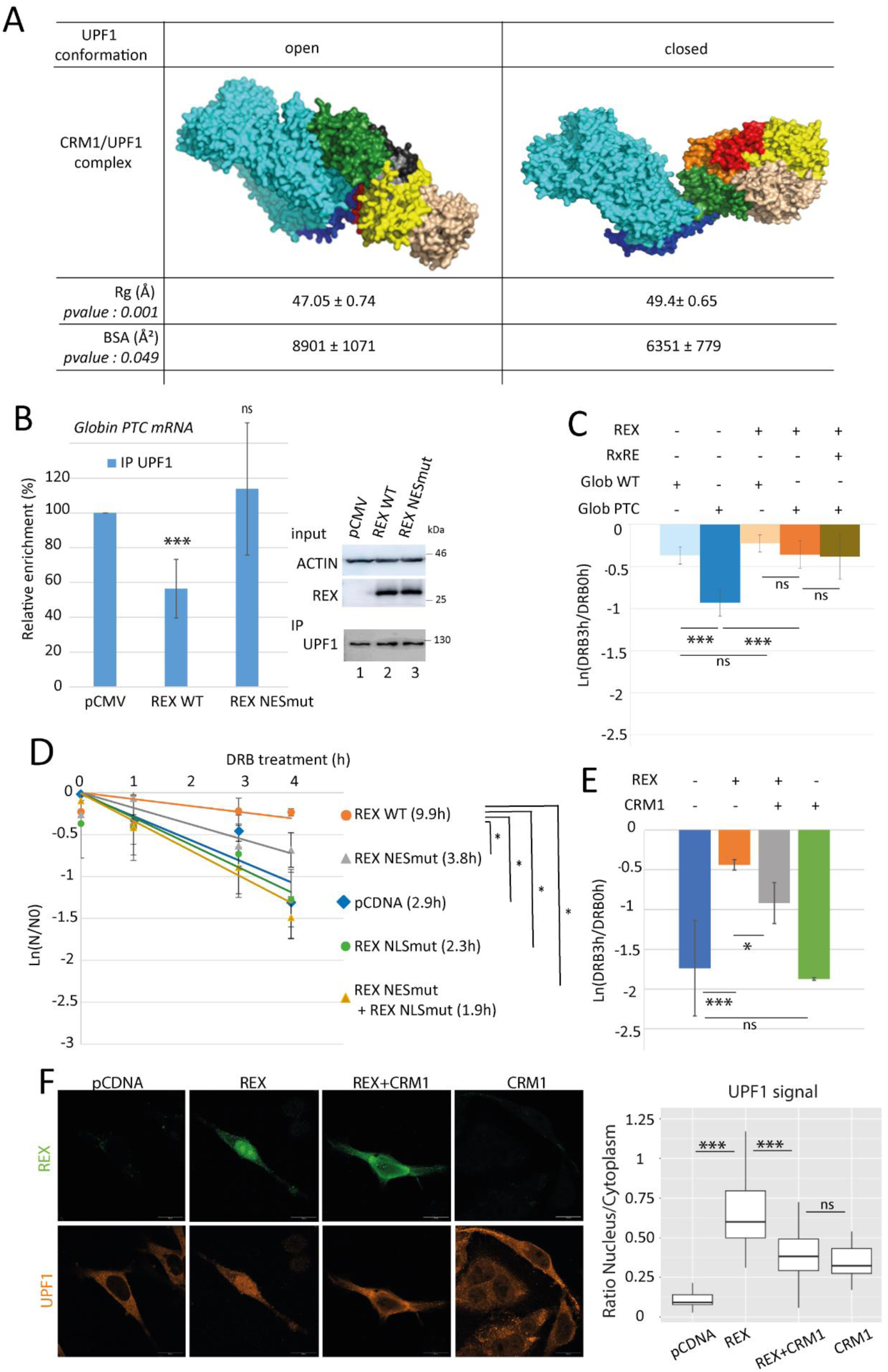
Rex expression leads to UPF1 reduced association to RNA, what is functionally link to NMD inhibition. A) Coarse-grained molecular modelling with Martini force field of CRM1-UPF1 complex starting from UPF1 which CH domain open (PDB 2WJV) of UPF1 with CH domain closed (PDB 2XZL). UPF1 fragment is 77-916. Molecular dynamics simulation using Gromacs with Martini 2 force-field during 1μs. UPF1 NES (89-103, dark blue) is bound to CRM1 (cyan) in both structures. UFP1 domains: CH: green; RecA2: wheat; RecA1: yellow, 1C: red, 1B: orange; stalk: black. The final state of simulations (1000ns) of UPF1-CRM1 complexes are presented. The complete simulation movie is available as an additional data. The “radius of gyration” (Rg(A)) and the “Buried Surface Area” (BSA (A²)) from 6 independent simulations are indicated. B) RNA immunoprecipitation experiments (RIP) using a rabbit polyclonal antibody targeting UPF1 were performed with HeLa cells transfected with a Globin PTC coding plasmid and WT or NES mutant Rex coding plasmid. Immunoprecipitated RNA (Globin PTC) were quantified by RTqPCR. For each condition, the relative enrichment of Globin PTC RNA associated to UPF1 compared to conditions without Rex was displayed in the graph. n=4 t-test pvalue: ns>0.05. and ***< 0.005). On the right panel, western blot controlling the levels of the expressed Rex and the immunoprecipitated UPF1. C) Decay rate analysis of Globin PTC and Globin WT RNA. Globin RNA levels were quantified after HeLa cells were treated with DRB for 0h or 3h. The rate of decay is expressed as ln(DRB3h/DRB0h). t-test pvalue: ns>0.05. and ***< 0.005. D) Half-life evaluation of the Glob PTC mRNA in HeLa cells transfected with the indicated forms of Rex. mRNA half-lifes (t1/2 = ln(2)/λ with λ the time constant of the decay curves) are indicated in front of their respective conditions. n=5 t-test pvalue: *< 0.05. The expression of Rex at each time point after DRB treatment was evaluated by WB (lower panel). E) Same as C with the Glob PTC mRNA and the indicated combinations of Rex and CRM1 overexpression. n=3 t-test pvalue: ns>0.05, *< 0.05 and ***<0.005. F) Confocal microscopy with Hela cells overexpressing the indicated combination of Rex and CRM1. UPF1 and Rex are stained. On the right, quantitative analysis of the microscopy experiments : UPF1 signal was quantified in the cytoplasm and in the nucleus and expressed as a ratio. Wilcoxon, pvalue ***< 0.005.

Nonsense Mediated mRNA Decay (NMD) is the most studied function of UPF1. As NMD requires UPF1 recruitment to RNA to initiate their degradation, we checked whether the defect of UPF1 binding on RNA upon Rex expression could correlate with NMD inhibition. To do so, we analysed the decay rate of the Globin PTC mRNA (that is an NMD sensitive reporter mRNA) with or without Rex expression, by monitoring the RNA levels after cell exposure to the transcription inhibitor DRB. Here, Rex expression led to a significant decrease in the decay rate of Globin PTC (**Fig 3C, compare dark blue and dark orange bars**). To control the specificity of this result, the decay rate of an NMD unsensitive RNA (Globin WT, a globin minigene without premature termination codon) was analysed as well. As expected, without Rex, the decay rate of Globin WT is lower than that of Globin PTC (**compare light and dark blue bars**). When Rex is expressed, the decay rate of Globin WT is unaffected (**compare light blue and orange bars)** and is of similar value as that of Globin PTC (**compare light and dark orange bars)**. The expression of a RxRE containing RNA did not significantly modify Rex’ impact on the Globin PTC decay rate (**compare dark orange and brown bars**). Finally, by using the sunset assay, we confirmed that these observations are not related to a possible translation inhibition by Rex (**supplementary Fig5B**). These results confirm that Rex specifically and efficiently inhibits NMD. To correlate this Rex dependent NMD inhibition with our previous observations, we analysed the effect of the Rex NES mutant on Globin PTC mRNA half-life (as another indication of NMD efficiency). Consistently with previous results, Globin PTC half-life is significantly increased by the expression of Rex WT compared to control condition (9.9h *vs* 2.9h) reflecting NMD inhibition. Conversely, Rex NES partially lost its ability to inhibit NMD as showed by the restored Globin PTC half-life (3.8h). As a control, we also monitored the effect of Rex NLS mutant, that couldn’t either support NMD inhibition (2.3h) (**Fig 3D and supplementary Fig5C**). Finally, to confirm that Rex expression inhibits NMD via the alteration of CRM1 behaviour, we measured the decay rates and half-lives of Globin PTC with different combinations of Rex and CRM1 overexpression (**Fig 3E** and **supplementary Fig 5D-E**): while Rex stabilized Globin PTC mRNA, the combined overexpression of Rex and CMR1 partially restored mRNA decay. To the opposite, CRM1 overexpression without Rex did not modify Globin PTC decay rate. Confocal microscopy observations were also performed with these four conditions, revealing that the NMD partial rescue was correlated with the partial restoration of UPF1 cellular distribution, as supported by the quantification of the nucleo/cytoplasmic ratio. Interestingly, we also observed a modification of Rex localization toward the cytoplasm or at the nuclear periphery (**Fig 3F and supplementary Fig5F**).

Altogether, these results suggest that Rex by favouring the interaction between UPF1 and CRM1 in the nucleus may decrease UPF1 association with RNA and inhibits NMD.

### 3. CRM1 hijacking by another retrovirus also interferes with UPF1 cellular localization and its NMD function

CRM1 hijacking is a hallmark of several complex retroviruses(45). For instance, HIV-1 express Rev, a functional homolog of HTLV-1 Rex; although these proteins share low sequence homology, they both display similar functional domains such as RNA binding domain, NLS, dual multimerization domain and NES. Rev also shuttles between nucleus and cytoplasm, hijacks CRM1 and exports the unspliced viral RNA (vRNA)(61). Moreover, a UPF1/Rev/CRM1 complex has already been described(62). As an alternative method to reinforce our observations that support an impact of CRM1 hijacking on UPF1 activity, we investigated the impact of Rev expression as we did previously with Rex. Thus, we directly looked at the endogenous UPF1 intracellular distribution under Rev expression by confocal microscopy. Rev strongly colocalizes with UPF1 and CRM1 in nucleolar compartments (**Fig. 4A (i) and supplementary Fig6A**). As for Rex, a significant nuclear retention of UPF1 can be quantified when Rev is expressed compared to control condition (**Fig 4A (ii)**). To further compare Rex and Rev impact on UPF1, we checked whether the Rev-mediated nuclear localisation of UPF1 can be also correlated with NMD inhibition. We found an increased half-life of Globin PTC after Rev expression compared to control condition (4.3h *vs* 1.6h, **Fig 4B**) suggesting that Rev is able to inhibit NMD and to alter the stability of non-viral RNAs. Moreover, as Rev-mediated inhibition of NMD is reported here for the first time, we further characterized the specificity of our observations: first, we excluded a translation defect by carrying out a sunset assay, as previously done for Rex (**supplementary Fig6B**). Second, we monitored Rev impact on the decay rate of Globin WT and compared it to Globin PTC. As expected, the decay rate of Globin WT was barely affected by Rev expression while that of Globin PTC was significantly decreased (**Fig 4C**). Furthermore, to determine whether these observations could be verified in the context of HIV-1 infection, we analyzed HeLa cells pre-transfected with Globin reporters and infected for 48h with HIV-GFP lentiviral particles (HIV) or lentiviral particles expressing GFP only (VLPCtr) as control. While the decay rate of Globin WT was not affected by HIV infection, that of Globin PTC was significantly reduced, indicating that HIV infection is able to inhibit NMD **(Fig 4D**). Altogether, these results demonstrate a specific NMD inhibition during HIV infection, which occurs in a Rev-dependent manner.

**Figure 4:**
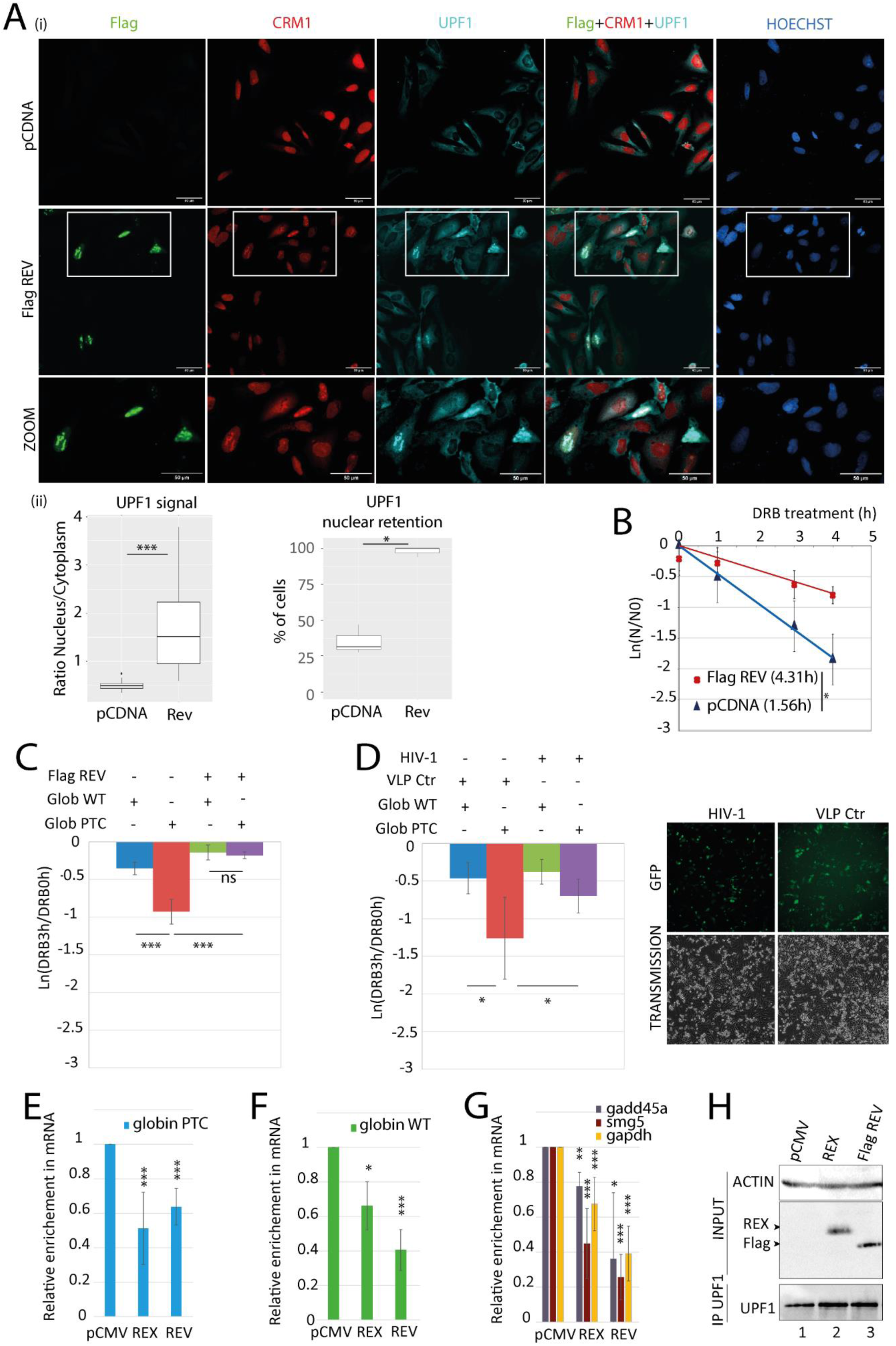
HIV-1 Rev promotes similar defects in UPF1 functions as HTLV-1 Rex. A) (i) Confocal microscopy experiments were performed in HeLa cells transfected with the plasmids indicated on the left side. Rev was revealed with an anti Flag tag antibody. Magnification of the framed zone is shown in the lower panel (ZOOM). Objective x20. Scale bar: 50µm. (ii) The same quantitative analysis was carried out as for Rex in Fig 2C. B) Half-life evaluation of the Glob PTC mRNA in HeLa cells transfected with the indicated forms of Rev. mRNA half-life (t1/2 = ln(2)/λ with λ the time constant of the decay curves) are indicated in front of their respective conditions. n=3 t-test * p<0.05. C) Decay rate analysis of Globin PTC and Globin WT RNA. Globin RNA levels were quantified after HeLa cells were treated with DRB for 0h or 3h. The rate of decay is expressed as ln (DRB3h/DRB0h). n=3 t-test ns: p>0.05; *** p<0.005. D) Same as C) except that HeLa cells were infected with HIV particles or control Virus Like Particles (CtrVLP) instead of being transfected with Flag Rev (left panel). GFP expressed from HIV or CtrVLP was observed with an epifluorescence microscope (right panel).). n=4 t-test * p<0.05 E) RIP using a rabbit polyclonal antibody targeting UPF1 was performed with HeLa cells expressing Globin PTC mRNA as well as Rex or Rev as indicated. Immunoprecipitated RNA (Globin PTC) were quantified by RTqPCR. For each condition, the relative enrichment of Globin PTC RNA associated to UPF1 compared to control conditions (pCMV) was displayed in the graph. n=4 t-test ns: p>0.05; * p<0.05, ** p<0.01, *** p<0.005 F) Same as E with cells expressing globin WT mRNA. G) same as E with the quantification of endogenous mRNA gadd45a, smg5 and gapdh. H) Representative western blot controlling the RIP experiments: Rex and Rev proteins expression was monitored in the whole cell extract (INPUT) as well as UPF1 in immunoprecipitations.

Consistently with our previous results on Rex, UPF1 RIP experiments revealed that Rev expression is also associated with a decrease in UPF1 association with the Globin PTC and the Globin WT mRNAs (**Fig 4E and 4F respectively**). To reinforce these observations, the association of UPF1 with endogenous NMD sensitive (*smg5* and *gadd45a*) and NMD unsensitive (*gapdh*) RNAs was also investigated and we observed a decrease in UPF1 association with these endogenous RNAs as well with Rev and Rex (**Fig 4G-H**). Therefore, Rev, like Rex, decreases/prevents UPF1 binding to cellular RNA, independently of their NMD sensitivity.

### 4. CRM1 hijacking by Rex in HTLV-1 transformed cell line affects UPF1 localisation and function

So far, we demonstrated that, independently of vRNA, Rex induced the nuclear retention of UPF1. We correlated it with higher levels of UPF1-CRM1 interaction, a reduced affinity for cellular RNA and NMD inhibition. Now, in order to investigate this process in physiologically relevant cellular models, we used three HTLV-1 chronically infected CD4+ T cell lines: C91PL and HUT102 that express Rex protein, and C8166 that displays a defect in Rex expression, as confirmed by western blot and confocal microscopy (**Fig 5A** and **supplementary Fig7B**). As a negative control we also used uninfected Jurkat CD4+ T cells. CoIP experiments confirm that UPF1 interacts with Rex in infected lymphocytes; moreover, in Rex expressing cells, UPF1 interacts with CRM1 in higher proportions than in Rex deprived cells (**Fig 5A, compare lanes 1-2 to 3-4**). Then, we examined the intracellular distribution of UPF1 and CRM1 in C91PL and C8166 by confocal microscopy. We found a nuclear retention of UPF1 in ∼75% of C91PL compared to ∼30% in Rex negative C8166 cells (**Fig 5B and supplementary Fig7A**). Z-stack analysis confirmed these conclusions (**Fig 5C**). Noteworthy, overlapping perinuclear foci of UPF1 and CRM1 were also observed in C91PL (**Fig 5B**). In the context of HTLV infection, the direct correlation between intracellular redistribution of UPF1/CRM1 and NMD inhibition is made difficult because both C8166 and C91PL cells express the viral protein Tax, another NMD inhibitor that we characterized in the past(38, 42). In these cell lines, the expression of Tax is well documented (38, 63) and its localization is not linked to either Rex or CRM1 distribution profiles (**supplementary Fig7B-C** and (64)). To evaluate NMD activity in a close model without Tax interference, we measured the half-life of the Globin PTC reporter in HeLa cells transfected with different mutated versions of a HTLV molecular clone (**supplementary Fig7D**). As expected, Rex expression alone could inhibit NMD. Finally, RIP showed a reduced enrichment of endogenous RNA (NMD sensitive or not) with UPF1 in C91PL compared to C8166 (**Fig 5D and supplementary Fig7E**). Altogether, these data corroborate our previous conclusions in a more physiological cellular model underlining their relevance (**Fig 5E**).

**Figure 5:**
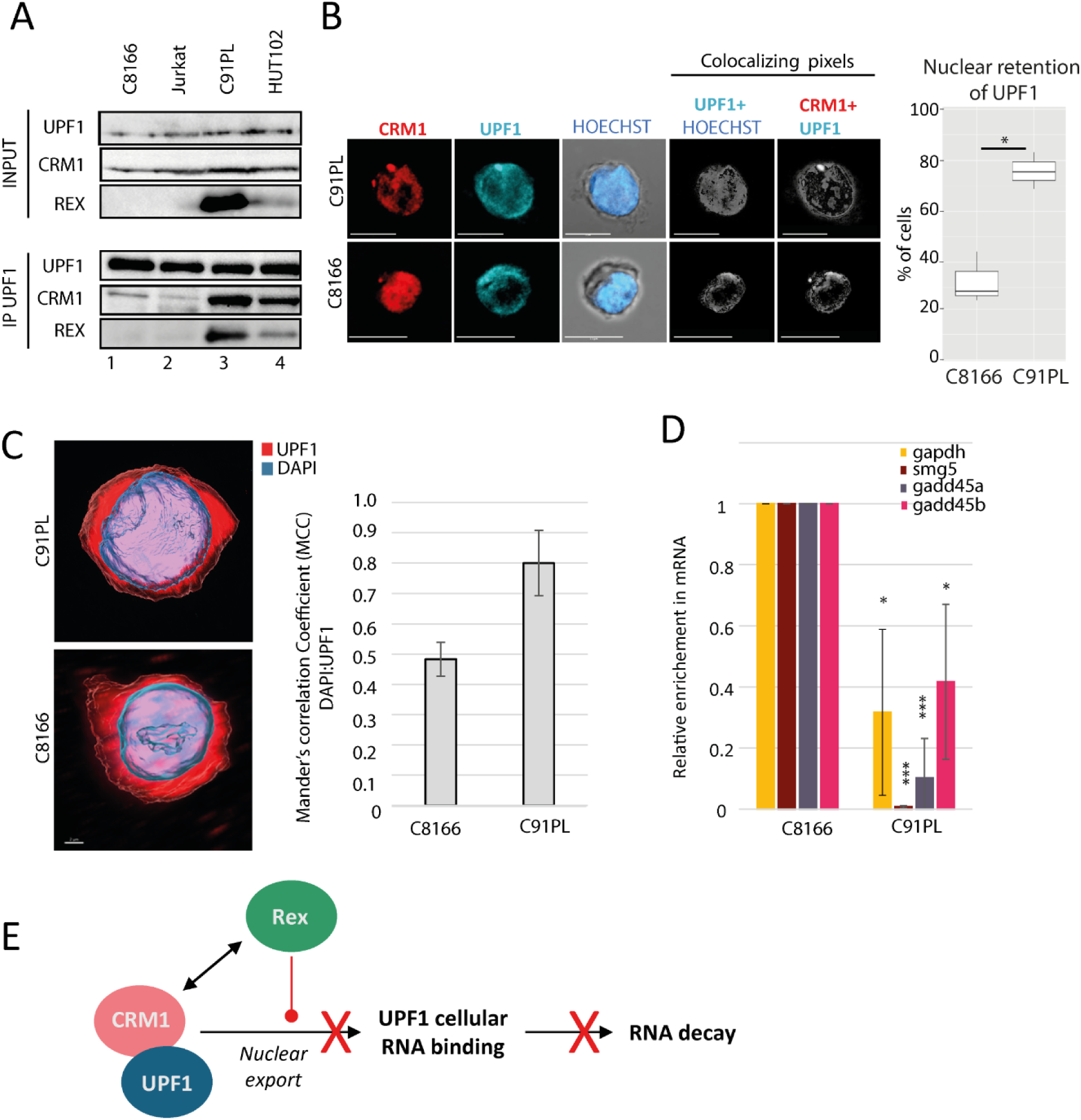
CRM1 hijacking by Rex affects UPF1 functions in infected lymphocytes. A) coIP experiments in the indicated lymphocytes cells extracts. IP were performed using rabbit polyclonal antibodies against UPF1. Proteins revealed by western blot are indicated on the side. B) (i) Confocal microscopy experiments were performed in C91PL and C8166 lymphocytes. CRM1 and UPF1 were revealed. Objective x63. Scale bar: 10µm. (ii) Quantitative analysis of UPF1 nuclear retention from confocal images C) Z stack analysis performed on C91PL and C8166 with a staining of the nucleus by DAPI and UPF1. The nuclear localisation of UPF1 is evaluated with the Mander’s correlation coefficient (MCC). Cells were obtained from 3 independent experiments. t-test pvalue: *<0.05. D) RIP using a rabbit polyclonal antibody targeting UPF1 were performed with C8166 and C91PL lymphocytes. Immunoprecipitated endogenous RNA(sensitive or not to NMD) were quantified by RTqPCR. The relative enrichment of RNA associated to UPF1 in C8166 compared to C91PL was displayed in the graph. n=3 t-test pvalue: *<0.05; ***<0.005. E) Graphical abstract of the consequences of CRM1 hijacking by Rex on the RNA helicase UPF1: Rex alters the nuclear export and favour the nuclear retention of NES containing cargoes, including UPF1. Concomitantly, Rex stabilizes CRM1/UPF1 association. Altogether, this lead to a defect in RNA binding and NMD inhibition.

### 5. UPF1 is addressed to vRNA in a Rex dependent manner

Unlike HeLa and 293T cells, C91PL cells constitutively express viral RNA. Knowing that the Rex/CRM1 complex binds selectively to unspliced viral RNA (vRNA) to drive its nuclear export, and that Rex and CRM1 modify UPF1 localisation and affinity for RNA, we questioned the interplay between UPF1 and vRNA. First, we carried out UPF1 RIP experiments in C8166 and C91PL cells, as well as in Jurkat T cells as a negative control. Unexpectedly, we found that UPF1 RIP were specifically enriched in vRNA in the Rex expressing cells only (**Fig 6A, supplementary Fig7F**). To investigate whether this association is related to NMD activation, we carried out RIP targeting UPF2, a NMD cofactor of UPF1. We found no enrichment of vRNA after UPF2 RIP in C91PL compared to C8166, in contrast to the RIP targeting UPF1 and Rex (**Fig 6B, supplementary Fig7G**). In addition, we performed UPF2 coIP in C91PL extracts: an interaction between UPF2 and CRM1 was found as expected since UPF2 is a known cargo of CRM1. However, no specific association between Rex and UPF2 could be found, unlike UPF1 (**Fig 6C**). Thus, these experiments demonstrate that the increased recruitment of UPF1 to vRNA and its incorporation in the vRNP is independent of UPF2 and subsequently independent of NMD activation. Then, we asked whether UPF1 and Rex can bind concomitantly to vRNA. To do so, we carried out double-RIP experiments. First, a RIP targeting a transiently expressed HA-tagged UPF1 was performed (**Fig 6D, blue bar**). Cells without HA-UPF1 expression were used as a control. Second, RNAs were natively eluted and further subjected to a second RIP against Rex (**Fig 6D, green bar**) or using a control immunoglobulin. RTqPCR on each RIP revealed a significant enrichment of vRNA associated to UPF1 as well as Rex, but to a lower amount. Thus, UPF1 can be recruited on vRNA concomitantly with Rex, in a Rex-dependent manner but independently of its NMD function. To clarify if this recruitment is controlled by the formation of the previously characterized CRM1-UPF1-Rex complex, RIP experiments were carried out on 293T cells transfected with the HTLV-R23L molecular clone (deficient for Rex expression) complemented with plasmids encoding Rex WT or Rex NES mutant. As a reminder, we have shown previously that Rex NES mutant can neither interact with CRM1 nor stabilize UPF1/CRM1 interaction (**Fig 2E**). RIP Rex experiments showed similar enrichment of vRNA with Rex WT and NES mutant proteins. In contrast, enrichment of vRNA in the UPF1 RIP was drastically reduced with the Rex NES mutant compared with the Rex WT protein (**Fig 6E**). Finally, we performed a RIP with UPF1 WT or NES mutant in 293T cells expressing a WT viral molecular clone: UPF1 NES mutant couldn’t bind vRNA to the contrary of UPF1 WT (**Fig 6F and supplementary Fig7H**). Altogether, these results indicate that in an infectious context, Rex orchestrates a CRM1 dependent switch in UPF1 RNA binding affinity: it drives the selective loading of UPF1 onto vRNA independently of the NMD process.

**Figure 6:**
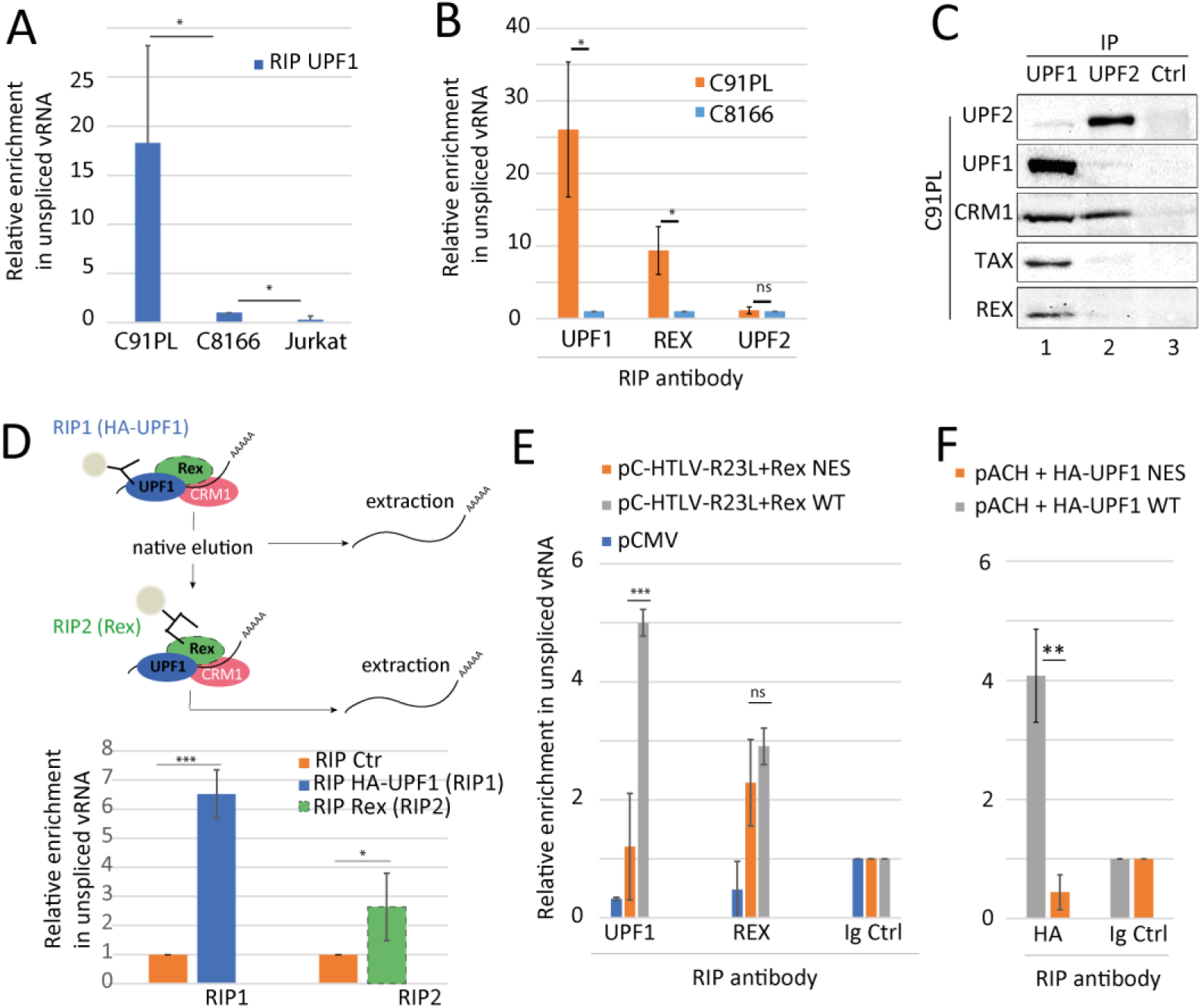
Rex/CRM1/UPF1 complex drives the loading of UPF1 on vRNA. A) RIP using a rabbit polyclonal antibody targeting UPF1 were performed with Jurkat, C8166 or C91PL lymphocytes. Immunoprecipitated unspliced viral RNA (vRNA) was quantified by RTqPCR. The relative enrichment of vRNA associated to UPF1 in Jurkat and C8166 compared to C91PL was displayed in the graph. n=5 t-test *** p<0.005 B) RIP experiments were carried out with the immunoprecipitation of UPF1, Rex or UPF2 as indicated in C91PL and in C8166 cells. The relative enrichment in vRNA in C91PL compared to C8166 is represented for each immunoprecipitated protein. n=3 t-test * p<0.05 C) coIP experiments in the C91PL cells extracts. IP were performed from the same extract using UPF1, UPF2 or a control antibody (Ctr). Proteins revealed by western blot were indicated on the side. D) Schematic diagram of the double RIP experiment performed in 293T cells co-transfected with the HTLV-1 WT molecular clone and a HA-UPF1 coding plasmid (upper panel). Quantification of the relative enrichment in vRNA after HA RIP targeting UPF1 (RIP1) and Rex RIP (RIP2) (lower panel) compared to RIP Ctr. n=3 t-test * p<0.05 E) RIP targeting UPF1, Rex or using a control antibody (as indicated) were performed with 293T cells transfected with HTLV-1 R23L molecular clone complemented with either Rex NES or Rex WT (orange and grey bars respectively). As a negative control, cells were transfected with an empty vector (blue bars). Immunoprecipitated RNA (vRNA) was quantified by RTqPCR. The graph displays the relative enrichment of vRNA associated to UPF1 or Rex compared to control antibody. n=4 t-test ns p>0.05, *** p<0.005. F) 293T cells are transfected with the indicated combinations of HTLV-1 molecular clone (pACH) and HA-UPF1 either WT (grey bars) or NES mutated (UPF1 NES, orange bars). Quantification of the relative enrichment in vRNA after RIP was performed with HA antibody or Ctr antibody. n=3 t-test ** p<0.01.

### 6. UPF1 supports different steps of HTLV-1 replication cycle leading to the production of viral particles

In this context, we hypothesized that the relocalization of UPF1 on vRNA induced by Rex and CRM1 might have a proviral function, contrasting with its antiviral function exerted via NMD. First, we investigated vRNA export. To do so, 293T cells were transduced with lentivectors expressing a doxycycline inducible UPF1 shRNA or a control shRNA. Cells were then treated twice with doxycycline to ensure the lowest levels of UPF1, prior to expression of Rex and a Luciferase reporter to monitor Rex-dependent export (FLucRxRE)(58). As shown in **Fig 7A**, in control conditions, Rex is required for the correct export of FLucRxRE RNA, resulting in firefly luciferase expression. To the contrary, the knock-down of UPF1 was associated with a decrease in firefly luciferase expression. To further link the hijacking of UPF1 in the vRNP and the stimulation of vRNA export, we test the impact of UPF1 NES mutant with this assay since we showed that this mutant is deficient in vRNA binding, even in the presence of Rex: first endogenous UPF1 levels are reduced with siRNA treatments to reduce the background; then different combinations of Rex and UPF1 WT or mutant plasmids are transfected. As expected, we found that UPF1 NES mutant has a reduced capacity to stimulate vRNA export, supporting the importance of the UPF1/CRM1 interaction and UPF1 loading on vRNA in the process (**Fig7B**). Accordingly, these results reveal a positive role for UPF1 in promoting Rex-dependent export of vRNA. Next, we evaluated the effects exerted by UPF1 during viral particles production steps. As previously mentioned, the fate of cytoplasmic vRNA is to be translated into structural viral proteins and enzymes or encapsidated in viral particles. To do so, we proceeded with UPF1 extinction before the expression of a replicative HTLV-1 molecular clone in 293T cells. Viral protein and vRNA from whole-cell extract or from the ECF (Extra Cellular Fraction that corresponds to cell-associated virions contained in viral biofilms) were analysed (**Fig 7C**). First, western blots showed that the levels of the GAG precursor, encoded by vRNA, were similar with or without UPF1(**Fig 7D (i)**). During viral particle formation, GAG maturation leads to its proteolytic cleavage into the nucleocapsid (NC/p15), the matrix (MA/p19) and the capsid (CA/p24). Unexpectedly, we observed by western blot that mature p19 and p24 were less present in whole-cell extracts deprived from UPF1 **(Fig 7D (i))**. To confirm this, FACS was performed on permeabilized cells using antibodies against UPF1 or p19. UPF1 knock-down was associated with a ∼50% decrease in the p19 mean fluorescence (**supplementary Fig8A**). We also performed ELISA on whole-cell extracts as well as on the corresponding ECF. In both fractions, UPF1 decrease was associated with weaker p19 signals (**Fig 7E, G**). Western blots confirmed that ECF from UPF1-depleted cells contained less p19 and p24. However, western blot on the ECF showed that the levels of viral envelope (Env) were less, if at all, affected by UPF1 knock-down **(Fig 7D(ii))**. Corroborating that observation, FACS analysis performed on non-permeabilized cells using the anti-HTLV serum or an anti-Env antibody did not reveal a significant decrease of the cell surface mean fluorescence **(supplementary Fig8B**). Concerning vRNA, qRT-PCR showed an increase in whole-cell extracts deprived from UPF1 that was expected, due to NMD inhibition. To the contrary, qRT-PCR on ECF purified RNA revealed a lower level of vRNA in the absence of UPF1 **(Fig 7F, H**). We previously demonstrated that the hijacking of UPF1 by Rex was linked with NMD inhibition. Thus, we checked whether the defect in p19 levels is directly due to the absence of UPF1 or indirectly due to NMD inhibition. Prior infection, 293T cells were treated with siRNA against UPF1 or UPF2: the monitoring of p19 by FACS confirmed that the absence of UPF1 provoked the reduction in p19 levels. To the contrary, UPF2 knock-down exerted no significant effect. The treatment of infected cells with the NMD inhibiting drug SMG1i, although strikingly preventing UPF1 phosphorylation necessary for NMD, had no significant effect on p19 either (**Fig 7I and supplementary Fig8C**). The data thus confirmed that the impact of UPF1 on GAG maturation is independent of NMD.

**Figure 7:**
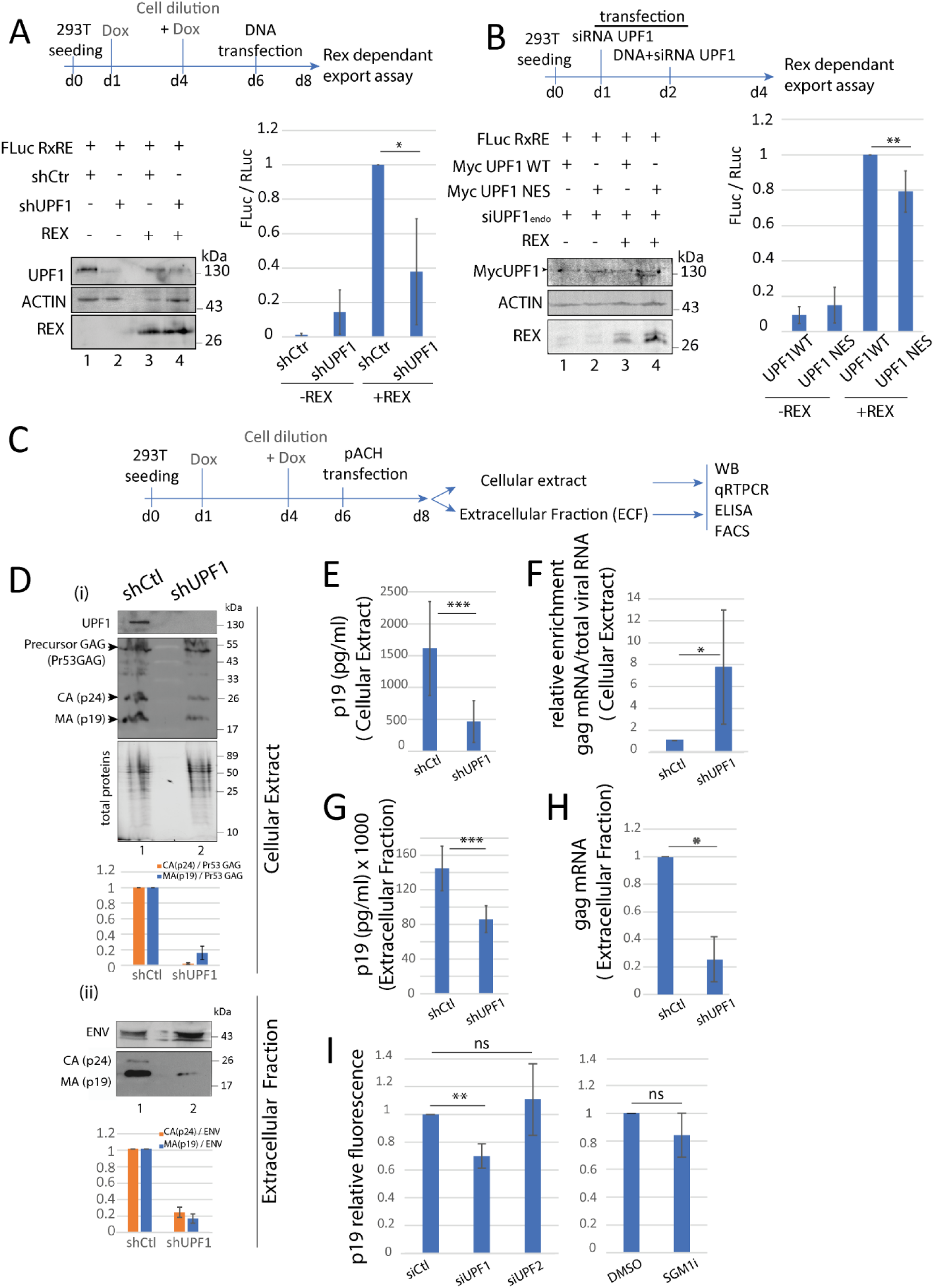
UPF1 controls different steps of the viral replication cycle. A) Protein expression in cell extracts used for dual luciferase assay was monitored by western blot (left). Dual luciferase assay was performed on shCtr or shUPF1 293T cells transfected with a Rex WT plasmid or an empty vector. Firefly luciferase as well as Renilla luciferase were quantified. Firefly (FLuc)/Renilla (RLuc) ratio was plotted in a graph. n=4 t-test * p<0.05 (right). B) Same as A with overexpression of MYC-UPF1 WT or mutated on the NES (MYC-UPF1 NES). To reduce the background of endogenous UPF1, cells were also treated with siRNA. Both Myc-UPF1 are siRNA resistant. C) Schematic representation of the experiment. The same preparation of cell extracts and the corresponding extracellular fractions were used to perform western blot, qRTPCR and ELISA. FACS data are presented in Supplementary Fig 5. D) Western blot was carried out on cellular extract (i) and on the corresponding extracellular fraction (ii). The recognized proteins were indicated on the left. The relative levels of the capsid (CA) and the matrix (MA) compared to the precursor GAG (i) or Env(ii) were evaluated after densitometry quantification using Imagelab software. E) ELISA spot assays (Zeptometrix) were performed with the cell extract of the indicated cells. MA(p19) quantification was plotted in bar plot. F) Same as E with the extracellular fraction preparation. G) vRNA quantifications were performed by qRTPCR on cell extracts. Primers used were localized in the gag coding region. n=5 t-test * p<0.05; ** p<0.01 H) Same as G with the corresponding ECF preparation. I) FACS quantifications of p19 protein in 293T cells treated with Ctr, UPF1 or UPF2 siRNA (left). The same experiment was performed with DMSO or the NMD inhibiting compound known as SMG1i. n=5 t-test ns p>0.05, ** p<0.01

Having demonstrated the loading of UPF1 on viral RNA, as well as the defect in vRNA packaging and viral particle maturation in the absence of UPF1, we wondered whether this might reflect the presence of UPF1 in the viral particles, where GAG maturation would occur after budding. First, we monitored the localisation of the viral matrix (MA/p19) and UPF1 in infected T cells. As expected, C8166 cannot express p19 due to a defect in Rex expression. In C91PL, p19 is displayed as foci at the cell periphery corresponding to the accumulation of virions on the cell surface(53). The immunostaining profiles of UPF1 and p19 IF overlap at these foci (**Fig 8A and supplementary Fig9A**). This suggests that UPF1 is present at budding sites and could effectively be incorporated into viral particles. To address this point, we isolated the ECF of 15×10^6^ infected cells, which contains most of the viral particles produced by the cell, forming a viral biofilm bound to the cell membrane as described previously(53, 65). The presence of viral particles was detected in C91PL only, as shown by monitoring p19 using western blot (**Fig 8B**). The absence of cellular contamination of ECF was controlled by probing the cytoplasmic protein RRM2. Finally, UPF1 was clearly identified in ECF from C91PL, but not on that of C8166 or controlled non-infected cells (**Fig 8B**). UPF1 RIP were also performed and revealed that UPF1 is bound to vRNA in ECF (**Fig 8C**). To quantitatively evaluate the levels of UPF1 in viral particles produced by C91PL infected cells, mass spectrometry experiments were performed on C91PL ECF as well as on purified virions released in C91PL culture supernatants (named"free particles"). UPF1 was detected in both fractions. As expected, UPF2 and RRM2 cannot be detected (**supplementary Fig9B**). The same experiments was also performed with uninfected CEM T-cells known to produce and released membrane bound extracellular vesicles (66, 67). A quantitative analysis showed that although UPF1 is present in the fraction derived from uninfected cells, it was significantly enriched in the C91PL one, that specifically contains viral particles **(Fig 8D and supplementary Table)**. However, UPF1 is identified as a relatively low concentrated protein in the HTLV-1 viral particles (**supplementary Fig 9C**). Altogether, these data confirm the presence of vRNA bound UPF1 in viral particles.

**Figure 8:**
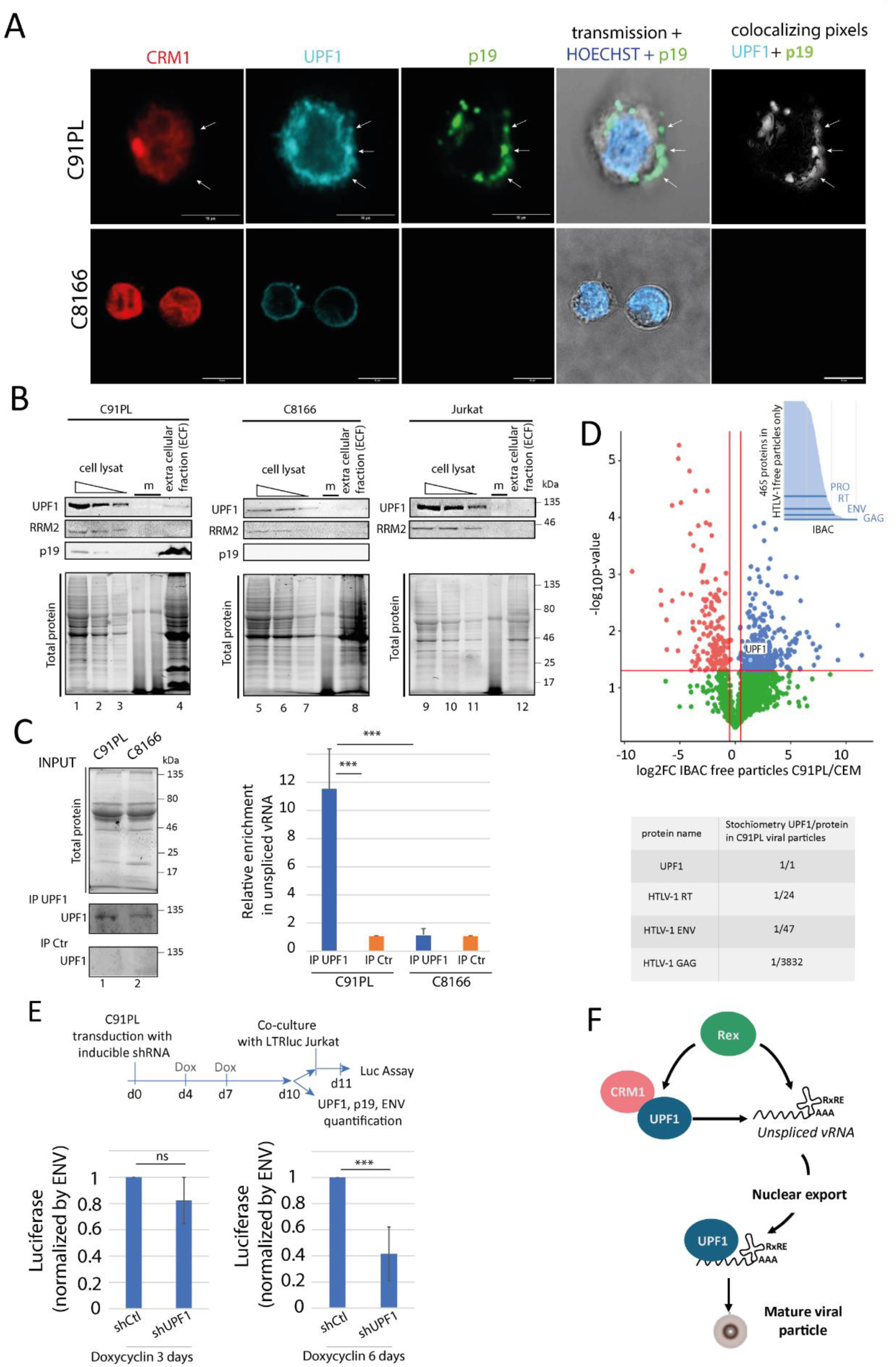
UPF1 is loaded into the viral particles produced by the virus. A) Confocal microscopy experiments were performed in C91PL and C8166 lymphocytes.: CRM1, UPF1 and p19 were revealed by IF. White arrows show p19 foci. Objective x63. Scale bar: 10µm. B) Extracellular fraction (ECF) containing the viral particles was collected from 15.10^6^ of C91PL, C8166 and Jurkat cells and analysed (lanes 4, 8, 12). The presence of UPF1, RRM2 and p19 was evaluated by WB and total protein was monitored with stainless procedure (upper and lower panels respectively). Cells used to harvest the ECF were also lysed and monitored by WB after serial dilutions (equivalent to 4.10^6^, 2.10^6^ and 10^6^ cells; lanes 1, 5, 9; 2, 6, 10; 3, 7, 11 respectively). C) RIP UPF1 on the ECF of C91PL and C8166. Western blot controls of the RIP on the left. Immunoprecipitated unspliced viral RNA (vRNA) was quantified by RTqPCR. The relative enrichment of vRNA associated to UPF1 compared to the control RIP was displayed in the graph on the right. n=3 t-test: *** p<0.005. D) Volcano plot of the log2FC (C91PL/CEM) for the “free particle” fraction. UPF1 enrichment in the C91PL samples compared to CEM samples is indicated (n=3). At the top, the distribution of the proteins only found in the C91PL samples is represented by their IBAC values: viral proteins are indicated. The stochiometric proportion of UPF1 compared to viral proteins from C91PL samples is also indicated (down). E) upper panel: schematic representation of the viral transmission experiment. Lower panel: bar plot representing the luciferase levels expressed by 150×10^3^ infected Jurkat after 24h of co-culture with 50×10^3^ C91PL. C91PL were treated with doxycyclin for 3 days (1 treatment) or 6 days (2 treatments) in order to express the shRNA (control or against UPF1). n=6, t-test ns p>0.05, *** p<0.005. F) Graphical abstract of the consequences of CRM1 hijacking by Rex on UPF1 interplay with vRNA.

Finally, we evaluated the impact of these modifications on HTLV viral particles infectivity and transmission. HTLV-1 transmission occurs via cell to cell contact, through viral biofilm in which viral particles are accumulated, and drastically improve the transmission rate(53). To assess the impact of UPF1 depletion on HTLV infectivity and spread, co-cultures between C91PL (expressing or not a UPF1 shRNA) and Jurkat-Luc reporter cells (expressing luciferase under the HTLV LTR) were performed as previously described(53). The significant reduction of UPF1, obtained after 6 days of shRNA induction, was associated with a decrease in p19 levels, consistent with our results on HTLV infected 293T cells (**supplementary Fig 9D**). Under these conditions, the decrease in UPF1 levels resulted in a drastic reduction (∼ 60%) of viral transmission to uninfected Jurkat cells after 24 hours of co-culture (**Fig 8E**). Collectively, these results showed that UPF1 defect altered the packaging of vRNA and the assembly or maturation of HTLV-1 viral particles. Ultimately, this multistep process support the virus capacity to transmit infectious virions to neighbouring cells.

In conclusion, we demonstrated that the hijacking of the exportin CRM1 by retroviral adapter such as HTLV-1 Rex, is combined with the hijack of the RNA helicase UPF1. On one side it affects cellular functions such as NMD; on the other side it favours the loading of UPF1 on vRNA to exerts functions independent of NMD, that promote viral replication and transmission (**Fig 8F**).

## Discussion

By regulating the intracellular distribution of hundreds of proteins and some specific RNA species, CRM1 export pathway is an important process controlling the cellular homeostasis. As introduced earlier, complex retroviruses like HTLV-1 and HIV-1 use retroviral adapters, respectively Rex and Rev, to export their viral RNA via CRM1. While the mechanistic steps of this export are already described, the cellular consequences of CRM1 hijacking have not been addressed yet. Here, we sought to characterize how the hijacking of CRM1 by Rex could affect the CRM1 cargo named UPF1, a RNA/DNA helicase involved in NMD, a well described RNA decay process with antiviral functions.

Our data showed that in Rex expressing cells, more CRM1 is associated to UPF1 especially in the nucleus. This complex relies on the combined interactions between UPF1, CRM1 and Rex. We hypothesized that we face a sequestration complex altering UPF1-CRM1 flux and recycling, ultimately repressing nuclear export. In this context, we confirmed that, UPF1 displays nuclear retention. Moreover, we proposed a model for the UPF1/CRM1 complex, based on the characterization of UPF1 NES domain and molecular dynamic simulations. Consistent with the observed enhanced interaction with CRM1, the most stable complex involves a conformation of UPF1 with a lower affinity for RNA(59). These simulations were challenged by RIP experiments *in cellulo* and we were able to link this UPF1-CRM1 complex with a defect in UPF1 binding to all cellular RNA, sensitive or not to NMD. Finally, we confirmed that NMD was repressed following Rex expression, as shown previously(39); more interestingly, we demonstrated that this inhibition can be due to the capacity of Rex to hijack and shuttle with CRM1. Similar results were obtained in HTLV-1 chronically infected cells, confirming their physiological relevance.

Recently, *Nakano et al* published that overexpressed UPF1 can interact with Rex in the cytoplasm and P-bodies and that the inhibition of NMD depends on several domains of Rex including its Nter part (ARM NLS and X domains)(43). Here, we report an interaction between Rex and endogenous UPF1, which is sensitive to a mutation in the NLS domain of Rex, in accordance with their observations. In addition, PLA experiments indicate that this interaction occurs in the cytoplasm as well as in the nucleus. Cytoplasmic complexes could match with the hypothesis of *Nakano et al*, who suggests that Rex is recruited to NMD complexes via UPF1, while nuclear foci would be relevant to the UPF1/CRM1/Rex complex that we described here and would support a model for UPF1 nuclear accumulation under Rex control. Indeed, using PLA, we also quantified an increase of nuclear UPF1/CRM1 foci upon Rex expression. In our model, to the contrary of *Nakano et al*, Rex ultimately prevents the incorporation of UPF1 in NMD complexes, as indicated by the lower affinity for RNA and the exclusion of UPF2. This inhibition is partially rescued when the cytoplasmic localisation of UPF1 is restored after overexpression of CRM1. This highlights that the Rex mediated NMD inhibition can rely on its ability to alter CRM1 export.

In order to reinforce the demonstration that CRM1 hijacking by a retroviral adapter can alter NMD, we alternatively investigated the effect of HIV-1 Rev: Rev is the functional homolog of Rex encoded by HIV-1 but both share little sequence identity. Although the sequential steps leading to the formation of an export complex with Rex and Rev may diverge, both result in the formation of a viral adapter multimer interacting with vRNA and CRM1. As expected, Rev expression led to similar observations, with slight differences: strikingly, Rev induced stronger UPF1 nuclear retention, essentially in nucleolus, while the nuclear staining of UPF1 was diffuse in the presence of Rex. These divergences may be due to specific mechanisms of CRM1 hijacking. Interestingly, this demonstration also points out for the first time that HIV-1 is able to trans-inhibit NMD in a Rev dependent manner, affecting the stability of non-viral RNA. In the cell, NMD exerts an important role during post-transcriptional regulation of gene expression. Notably, it controls pathways involved in the equilibrium between proliferation and differentiation, stress response and immune response. With regards to this role, it is expected that CRM1 hijacking by Rex and Rev lead to specific post-transcriptional deregulations due to NMD inhibition. Some of these deregulations have already been addressed(68) but a dedicated study is still needed to unravel the significance of these effects during viral infection.

During HIV-1 replication, UPF1 is involved in vRNA export with Rev and CRM1; moreover, in silico protein-protein docking analyses suggested that these interactions occur in a region that overlaps with the UPF2 binding site of UPF1, thereby protecting vRNA from NMD activation(62), (69). UPF1 does not affect the viral particle formation but is encapsidated and favors post-entry steps of the infection(70). With HTLV-1, we found that CRM1 ability to interact with Rex and UPF1 favours this latter recruitment onto vRNA. Considering our observations about UPF1-CRM1 complex affinity for RNA, it is probable that Rex mediates this recruitment via its RNA binding motif and the RxRE motif of vRNA. Thus, UPF1 undergoes a switch from cellular mRNA to vRNA that is not linked with NMD activation, as it occurs independently from UPF2. We clearly established that UPF1 plays a positive role in the Rex-dependent vRNA export. Furthermore, we found that in the absence of UPF1, the maturation of structural proteins MA(p19) and CA(p24) was impaired: a decrease in MA and CA levels was observed in cell extracts as well as in the extra-cellular fraction where infectious viral particles accumulate. This reduction is unlikely to be due to the repression of vRNA export, since the GAG precursor (pr53GAG) levels were barely affected. Moreover, NMD inhibition by UPF2 extinction or SMG1 inhibition didn’t affect MA maturation, confirming that UPF1 acts in an NMD independent manner. Importantly, the expression of the viral envelope was not significantly affected. Altogether, those observations revealed a role of UPF1 in establishing a correct balance between the different structural proteins that compose the viral particle. We may hypothesize that at the site of viral particles budding, UPF1, bound to vRNA, controls specific maturation steps leading to the production of complete, infectious virions. In support of this hypothesis, we provided evidence that (i) UPF1 is loaded itself into viral particles bound to vRNA and that in its absence, vRNA are less incorporated in the viral particles; (ii) viral infectivity from C91PL was clearly repressed when UPF1 levels were reduced. The importance of helicases during the packaging steps has been reported for some viruses in the past: Rep52/40 DNA helicases from AAV-2 are required to translocate single-stranded DNA genomes into preformed empty capsids(71) while NS3 is required for the packaging of YFV vRNA(72). Retroviruses that lack viral helicases rely on the host’s ones : DDX6 is re-directed to the assembly site of foamy virus particles and drags vRNA into them(73); for HIV-1, DDX6 directly facilitates capsid assembly while DDX24 promotes the packaging of viral RNA(74). The new function revealed in this work for UPF1 during HTLV-1 maturation might be in line with this. However, deciphering the underlying molecular mechanisms will require further investigation.

Finally, this work opens new perspectives on the regulation of NMD: in the past, the nuclear import of UPF1 by the importin β1 was stressed out as a sensitive step (75). Here, the expression of viral proteins shows that the modulation of the CRM1 dependent export also impacts on NMD, underscoring the importance of the nucleo/cytoplasmic shuttling process as a node of NMD regulation. In line with this, our study led to the precise delineation of the NES domain of UPF1 and a model of interaction between both proteins. Further investigations are needed to understand how the NMD can be controlled at this step in the absence of viral infection.

In conclusion, these results are the proof of concept that CRM1 subjugation by retroviral proteins can affect cellular pathways. Notably, it trans-inhibits the NMD via the formation of a complex trapping UPF1 in a conformation with low affinity for RNA. Unexpectedly, the hijacked UPF1 shifts from an antiviral to a proviral function: it is redirected on the vRNA with Rex, where it is isolated from its NMD partner UPF2 to stimulate vRNA export and viral particles maturation.

## Data availability

The data underlying this article are available in the article and in its online supplementary material. Mass spectrometry data are available via ProteomeXchange with identifier PXD043988. Reviewer account details: **Username:**;

The pipeline for the nucleus/cytoplasm analysis is available at https://gitbio.ens-lyon.fr/sdebosso/mocquet/

## Acknowledgments

We would like to thank Derse’s lab for the pCMV-HTLV-1 plasmids and A. Gessain for anti HTLV-1 serum. We also thank S. Alais for technical support with ELISA and the harvest of lymphocytes extracellular matrix, P. Solanes for help in setting up the proteomic study, F. Mortreux, C. Pique and C. Journo for useful comments on the manuscript. We acknowledge the contribution of SFR Biosciences (UMS3444/CNRS, US8/Inserm, ENS de Lyon, UCBL) facilities: the PLATIM and especially C. Lionnet and E. Chartre for the assistance with microscopy experiments; the center AniRA from CEPHALIA infrastructure (FACS and vectorology) and especially Gisèle Froment, Didier Nègre and C. Costa.

## Author Contributions statement

Investigation and formal analysis, L.P., M.MH, A.R., M.P., JP.R., L.S., S.D., D.C., M.M., D.D, M.M., S.R., A.D. and V.M.; Conceptualization, V.M.; writing, L.P., M.MH., H.D., MI.T., F.L., P.J. and V.M.; funding acquisition, M.MH, L.P., MI.T and V.M.

## References

1. Okamura, M., Inose, H. and Masuda, S. (2015) RNA Export through the NPC in Eukaryotes. Genes, 6, 124– 149.

2. Cautain, B., Hill, R., de Pedro, N. and Link, W. (2015) Components and regulation of nuclear transport processes. FEBS J., 282, 445–462.

3. Fung, H.Y.J., Niesman, A. and Chook, Y.M. (2021) An update to the CRM1 cargo/NES database NESdb. Mol. Biol. Cell, 32, 467–469.

4. Fung, H.Y.J. and Chook, Y.M. (2014) Atomic basis of CRM1-cargo recognition, release and inhibition. Semin. Cancer Biol., 27, 52–61.

5. Güttler, T. and Görlich, D. (2011) Ran-dependent nuclear export mediators: a structural perspective. EMBO J., 30, 3457–3474.

6. Güttler, T., Madl, T., Neumann, P., Deichsel, D., Corsini, L., Monecke, T., Ficner, R., Sattler, M. and Görlich, D. (2010) NES consensus redefined by structures of PKI-type and Rev-type nuclear export signals bound to CRM1. Nat. Struct. Mol. Biol., 17, 1367–1376.

7. Kehlenbach, R.H., Dickmanns, A. and Gerace, L. (1998) Nucleocytoplasmic shuttling factors including Ran and CRM1 mediate nuclear export of NFAT In vitro. J. Cell Biol., 141, 863–874.

8. Fu, S.-C., Fung, H.Y.J., Cağatay, T., Baumhardt, J. and Chook, Y.M. (2018) Correlation of CRM1-NES affinity with nuclear export activity. Mol. Biol. Cell, 29, 2037–2044.

9. Engelsma, D., Bernad, R., Calafat, J. and Fornerod, M. (2004) Supraphysiological nuclear export signals bind CRM1 independently of RanGTP and arrest at Nup358. EMBO J., 23, 3643–3652.

10. Zhang, X., Yamada, M., Mabuchi, N. and Shida, H. (2003) Cellular requirements for CRM1 import and export. J. Biochem. (Tokyo), 134, 759–764.

11. Hill, R., Cautain, B., de Pedro, N. and Link, W. (2014) Targeting nucleocytoplasmic transport in cancer therapy. Oncotarget, 5, 11–28.

12. Nguyen, K.T., Holloway, M.P. and Altura, R.A. (2012) The CRM1 nuclear export protein in normal development and disease. Int. J. Biochem. Mol. Biol., 3, 137–151.

13. Saulino, D.M., Younes, P.S., Bailey, J.M. and Younes, M. (2018) CRM1/XPO1 expression in pancreatic adenocarcinoma correlates with survivin expression and the proliferative activity. Oncotarget, 9, 21289–21295.

14. Chen, Y., Camacho, S.C., Silvers, T.R., Razak, A.R.A., Gabrail, N.Y., Gerecitano, J.F., Kalir, E., Pereira, E., Evans, B.R., Ramus, S.J., et al. (2017) Inhibition of the Nuclear Export Receptor XPO1 as a Therapeutic Target for Platinum-Resistant Ovarian Cancer. Clin. Cancer Res. Off. J. Am. Assoc. Cancer Res., 23, 1552–1563.

15. Jiang, Y., Hou, J., Zhang, X., Xu, G., Wang, Y., Shen, L., Wu, Y., Li, Y. and Yao, L. (2020) Circ-XPO1 upregulates XPO1 expression by sponging multiple miRNAs to facilitate osteosarcoma cell progression. Exp. Mol. Pathol., 117, 104553.

16. Song, P., Li, W., Xie, J., Hou, Y. and You, C. (2020) Cytokine storm induced by SARS-CoV-2. Clin. Chim. Acta Int. J. Clin. Chem., 509, 280–287.

17. Ferreira, B.I., Cautain, B., Grenho, I. and Link, W. (2020) Small Molecule Inhibitors of CRM1. Front. Pharmacol., 11, 625.

18. Wang, A.Y. and Liu, H. (2019) The past, present, and future of CRM1/XPO1 inhibitors. Stem Cell Investig., 6, 6.

19. Kırlı, K., Karaca, S., Dehne, H.J., Samwer, M., Pan, K.T., Lenz, C., Urlaub, H. and Görlich, D. (2015) A deep proteomics perspective on CRM1-mediated nuclear export and nucleocytoplasmic partitioning. eLife, 4, e11466.

20. Thakar, K., Karaca, S., Port, S.A., Urlaub, H. and Kehlenbach, R.H. (2013) Identification of CRM1-dependent Nuclear Export Cargos Using Quantitative Mass Spectrometry. Mol. Cell. Proteomics MCP, 12, 664– 678.

21. Mendell, J.T., Sharifi, N.A., Meyers, J.L., Martinez-Murillo, F. and Dietz, H.C. (2004) Nonsense surveillance regulates expression of diverse classes of mammalian transcripts and mutes genomic noise. Nat Genet, 36, 1073–8.

22. Lejeune, F. (2022) Nonsense-Mediated mRNA Decay, a Finely Regulated Mechanism. Biomedicines, 10, 141.

23. Hurt, J.A., Robertson, A.D. and Burge, C.B. (2013) Global analyses of UPF1 binding and function reveal expanded scope of nonsense-mediated mRNA decay. Genome Res, 23, 1636–1650.

24. Zund, D., Gruber, A.R., Zavolan, M. and Muhlemann, O. (2013) Translation-dependent displacement of UPF1 from coding sequences causes its enrichment in 3’ UTRs. Nat Struct Mol Biol, 20, 936–43.

25. Neu-Yilik, G., Raimondeau, E., Eliseev, B., Yeramala, L., Amthor, B., Deniaud, A., Huard, K., Kerschgens, K., Hentze, M.W., Schaffitzel, C., et al. (2017) Dual function of UPF3B in early and late translation termination. EMBO J., 36, 2968–2986.

26. Chamieh, H., Ballut, L., Bonneau, F. and Le Hir, H. (2008) NMD factors UPF2 and UPF3 bridge UPF1 to the exon junction complex and stimulate its RNA helicase activity. Nat Struct Mol Biol, 15, 85–93.

27. Yamashita, A., Ohnishi, T., Kashima, I., Taya, Y. and Ohno, S. (2001) Human SMG-1, a novel phosphatidylinositol 3-kinase-related protein kinase, associates with components of the mRNA surveillance complex and is involved in the regulation of nonsense-mediated mRNA decay. Genes Dev, 15, 2215–28.

28. Palma, M., Leroy, C., Salomé-Desnoulez, S., Werkmeister, E., Kong, R., Mongy, M., Le Hir, H. and Lejeune, F. (2021) A role for AKT1 in nonsense-mediated mRNA decay. Nucleic Acids Res., 49, 11022–11037.

29. Cho, H., Abshire, E.T., Popp, M.W., Pröschel, C., Schwartz, J.L., Yeo, G.W. and Maquat, L.E. (2022) AKT constitutes a signal-promoted alternative exon-junction complex that regulates nonsense-mediated mRNA decay. Mol. Cell, 82, 2779–2796.e10.

30. Durand, S., Franks, T.M. and Lykke-Andersen, J. (2016) Hyperphosphorylation amplifies UPF1 activity to resolve stalls in nonsense-mediated mRNA decay. Nat. Commun., 7, 12434.

31. Balistreri, G., Horvath, P., Schweingruber, C., Zünd, D., McInerney, G., Merits, A., Mühlemann, O., Azzalin, C. and Helenius, A. (2014) The host nonsense-mediated mRNA decay pathway restricts Mammalian RNA virus replication. Cell Host Microbe, 16, 403–411.

32. Hogg, J.R. and Goff, S.P. (2010) Upf1 senses 3’UTR length to potentiate mRNA decay. Cell, 143, 379–89.

33. LeBlanc, J.J. and Beemon, K.L. (2004) Unspliced Rous sarcoma virus genomic RNAs are translated and subjected to nonsense-mediated mRNA decay before packaging. J. Virol., 78, 5139–5146.

34. Wada, M., Lokugamage, K.G., Nakagawa, K., Narayanan, K. and Makino, S. (2018) Interplay between coronavirus, a cytoplasmic RNA virus, and nonsense-mediated mRNA decay pathway. Proc. Natl. Acad. Sci. U. S. A., 115, E10157–E10166.

35. Quek, B.L. and Beemon, K. (2014) Retroviral strategy to stabilize viral RNA. Curr. Opin. Microbiol., 18, 78– 82.

36. Garcia, D., Garcia, S. and Voinnet, O. (2014) Nonsense-mediated decay serves as a general viral restriction mechanism in plants. Cell Host Microbe, 16, 391–402.

37. Li, M., Johnson, J.R., Truong, B., Kim, G., Weinbren, N., Dittmar, M., Shah, P.S., Von Dollen, J., Newton, B.W., Jang, G.M., et al. (2019) Identification of antiviral roles for the exon-junction complex and nonsense-mediated decay in flaviviral infection. Nat. Microbiol., 4, 985–995.

38. Mocquet, V., Neusiedler, J., Rende, F., Cluet, D., Robin, J.P., Terme, J.M., Duc Dodon, M., Wittmann, J., Morris, C., Le Hir, H., et al. (2012) The human T-lymphotropic virus type 1 tax protein inhibits nonsense-mediated mRNA decay by interacting with INT6/EIF3E and UPF1. J Virol, 86, 7530–43.

39. Nakano, K., Ando, T., Yamagishi, M., Yokoyama, K., Ishida, T., Ohsugi, T., Tanaka, Y., Brighty, D.W. and Watanabe, T. (2013) Viral interference with host mRNA surveillance, the nonsense-mediated mRNA decay (NMD) pathway, through a new function of HTLV-1 Rex: implications for retroviral replication. Microbes Infect., 15, 491–505.

40. Popp, M.W.-L., Cho, H. and Maquat, L.E. (2020) Viral subversion of nonsense-mediated mRNA decay. RNA N. Y. N, 26, 1509–1518.

41. Nuccetelli, V., Mghezzi-Habellah, M., Deymier, S., Roisin, A., Gérard-Baraggia, F., Rocchi, C., Coureux, P.-D., Gouet, P., Cimarelli, A., Mocquet, V., et al. (2025) The SARS-CoV-2 nucleocapsid protein interferes with the full enzymatic activation of UPF1 and its interaction with UPF2. Nucleic Acids Res., 53, gkaf010.

42. Fiorini, F., Robin, J.-P., Kanaan, J., Borowiak, M., Croquette, V., Le Hir, H., Jalinot, P. and Mocquet, V. (2018) HTLV-1 Tax plugs and freezes UPF1 helicase leading to nonsense-mediated mRNA decay inhibition. Nat. Commun., 9.

43. Nakano, K., Karasawa, N., Hashizume, M., Tanaka, Y., Ohsugi, T., Uchimaru, K. and Watanabe, T. (2022) Elucidation of the Mechanism of Host NMD Suppression by HTLV-1 Rex: Dissection of Rex to Identify the NMD Inhibitory Domain. Viruses, 14, 344.

44. Bogerd, H.P., Echarri, A., Ross, T.M. and Cullen, B.R. (1998) Inhibition of human immunodeficiency virus Rev and human T-cell leukemia virus Rex function, but not Mason-Pfizer monkey virus constitutive transport element activity, by a mutant human nucleoporin targeted to Crm1. J. Virol., 72, 8627–8635.

45. Shida, H. (2012) Role of Nucleocytoplasmic RNA Transport during the Life Cycle of Retroviruses. Front. Microbiol., 3, 179.

46. Nakano, K. and Watanabe, T. (2012) HTLV-1 Rex: the courier of viral messages making use of the host vehicle. Front. Microbiol., 3, 330.

47. Ahmed, Y.F., Hanly, S.M., Malim, M.H., Cullen, B.R. and Greene, W.C. (1990) Structure-function analyses of the HTLV-I Rex and HIV-1 Rev RNA response elements: insights into the mechanism of Rex and Rev action. Genes Dev., 4, 1014–1022.

48. Palmeri, D. and Malim, M.H. (1999) Importin beta can mediate the nuclear import of an arginine-rich nuclear localization signal in the absence of importin alpha. Mol. Cell. Biol., 19, 1218–1225.

49. Ballaun, C., Farrington, G.K., Dobrovnik, M., Rusche, J., Hauber, J. and Böhnlein, E. (1991) Functional analysis of human T-cell leukemia virus type I rex-response element: direct RNA binding of Rex protein correlates with in vivo activity. J. Virol., 65, 4408–4413.

50. Hakata, Y., Umemoto, T., Matsushita, S. and Shida, H. (1998) Involvement of human CRM1 (exportin 1) in the export and multimerization of the Rex protein of human T-cell leukemia virus type 1. J. Virol., 72, 6602–6607.

51. Nakano, K. and Watanabe, T. (2016) HTLV-1 Rex Tunes the Cellular Environment Favorable for Viral Replication. Viruses, 8, 58.

52. Mazurov, D., Ilinskaya, A., Heidecker, G., Lloyd, P. and Derse, D. (2010) Quantitative comparison of HTLV-1 and HIV-1 cell-to-cell infection with new replication dependent vectors. PLoS Pathog., 6, e1000788.

53. Pais-Correia, A.-M., Sachse, M., Guadagnini, S., Robbiati, V., Lasserre, R., Gessain, A., Gout, O., Alcover, A. and Thoulouze, M.-I. (2010) Biofilm-like extracellular viral assemblies mediate HTLV-1 cell-to-cell transmission at virological synapses. Nat. Med., 16, 83–89.

54. Cox, J., Neuhauser, N., Michalski, A., Scheltema, R.A., Olsen, J.V. and Mann, M. (2011) Andromeda: a peptide search engine integrated into the MaxQuant environment. J. Proteome Res., 10, 1794–1805.

55. Tyanova, S., Temu, T. and Cox, J. (2016) The MaxQuant computational platform for mass spectrometry-based shotgun proteomics. Nat. Protoc., 11, 2301–2319.

56. Cachat, A., Alais, S., Chevalier, S.A., Journo, C., Fusil, F., Dutartre, H., Boniface, A., Ko, N.L., Gessain, A., Cosset, F.-L., et al. (2014) ADAR1 enhances HTLV-1 and HTLV-2 replication through inhibition of PKR activity. Retrovirology, 11, 93.

57. Niopek, D., Wehler, P., Roensch, J., Eils, R. and Di Ventura, B. (2016) Optogenetic control of nuclear protein export. Nat. Commun., 7, 10624.

58. Kress, E., Baydoun, H.H., Bex, F., Gazzolo, L. and Duc Dodon, M. (2005) Critical role of hnRNP A1 in HTLV-1 replication in human transformed T lymphocytes. Retrovirology, 2, 8.

59. Langer, L.M., Kurscheidt, K., Basquin, J., Bonneau, F., Iermak, I., Basquin, C. and Conti, E. (2024) UPF1 helicase orchestrates mutually exclusive interactions with the SMG6 endonuclease and UPF2. Nucleic Acids Res., 52, 6036–6048.

60. Thermann, R., Neu-Yilik, G., Deters, A., Frede, U., Wehr, K., Hagemeier, C., Hentze, M.W. and Kulozik, A.E. (1998) Binary specification of nonsense codons by splicing and cytoplasmic translation. Embo J, 17, 3484–94.

61. Behrens, R.T., Aligeti, M., Pocock, G.M., Higgins, C.A. and Sherer, N.M. (2017) Nuclear Export Signal Masking Regulates HIV-1 Rev Trafficking and Viral RNA Nuclear Export. J. Virol., 91, e02107–16.

62. Ajamian, L., Abel, K., Rao, S., Vyboh, K., García-de-Gracia, F., Soto-Rifo, R., Kulozik, A.E., Gehring, N.H. and Mouland, A.J. (2015) HIV-1 Recruits UPF1 but Excludes UPF2 to Promote Nucleocytoplasmic Export of the Genomic RNA. Biomolecules, 5, 2808–2839.

63. Liu, B., Hong, S., Tang, Z., Yu, H. and Giam, C.-Z. (2005) HTLV-I Tax directly binds the Cdc20-associated anaphase-promoting complex and activates it ahead of schedule. Proc. Natl. Acad. Sci. U. S. A., 102, 63–68.

64. Baydoun, H.H., Bellon, M. and Nicot, C. (2008) HTLV-1 Yin and Yang: Rex and p30 master regulators of viral mRNA trafficking. AIDS Rev., 10, 195–204.

65. Alais, S., Mahieux, R. and Dutartre, H. (2015) Viral Source-Independent High Susceptibility of Dendritic Cells to Human T-Cell Leukemia Virus Type 1 Infection Compared to That of T Lymphocytes. J. Virol., 89, 10580–10590.

66. Soo, C.Y., Song, Y., Zheng, Y., Campbell, E.C., Riches, A.C., Gunn-Moore, F. and Powis, S.J. (2012) Nanoparticle tracking analysis monitors microvesicle and exosome secretion from immune cells: Nanoparticle tracking analysis of microvesicles and exosomes. Immunology, 136, 192–197.

67. Liu, Y.-J. and Wang, C. (2023) A review of the regulatory mechanisms of extracellular vesicles-mediated intercellular communication. Cell Commun. Signal. CCS, 21, 77.

68. Nakano, K., Yokoyama, K., Shin, S., Uchida, K., Tsuji, K., Tanaka, M., Uchimaru, K. and Watanabe, T. (2022) Exploring New Functional Aspects of HTLV-1 RNA-Binding Protein Rex: How Does Rex Control Viral Replication? Viruses, 14, 407.

69. Ajamian, L., Abrahamyan, L., Milev, M., Ivanov, P.V., Kulozik, A.E., Gehring, N.H. and Mouland, A.J. (2008) Unexpected roles for UPF1 in HIV-1 RNA metabolism and translation. RNA N. Y. N, 14, 914–927.

70. Serquina, A.K., Das, S.R., Popova, E., Ojelabi, O.A., Roy, C.K. and Gottlinger, H.G. (2013) UPF1 Is Crucial for the Infectivity of Human Immunodeficiency Virus Type 1 Progeny Virions. J Virol, 87, 8853–61.

71. King, J.A., Dubielzig, R., Grimm, D. and Kleinschmidt, J.A. (2001) DNA helicase-mediated packaging of adeno-associated virus type 2 genomes into preformed capsids. EMBO J., 20, 3282–3291.

72. Patkar, C.G. and Kuhn, R.J. (2008) Yellow Fever virus NS3 plays an essential role in virus assembly independent of its known enzymatic functions. J. Virol., 82, 3342–3352.

73. Yu, S.F., Lujan, P., Jackson, D.L., Emerman, M. and Linial, M.L. (2011) The DEAD-box RNA Helicase DDX6 is Required for Efficient Encapsidation of a Retroviral Genome. PLoS Pathog., 7, e1002303.

74. Ma, J., Rong, L., Zhou, Y., Roy, B.B., Lu, J., Abrahamyan, L., Mouland, A.J., Pan, Q. and Liang, C. (2008) The requirement of the DEAD-box protein DDX24 for the packaging of human immunodeficiency virus type 1 RNA. Virology, 375, 253–264.

75. Hu, J., Li, P., Shi, B. and Tie, J. (2021) Importin β1 mediates nuclear import of the factors associated with nonsense-mediated RNA decay. Biochem. Biophys. Res. Commun., 542, 34–39.

