## Supplementary Figures for "Retroviral adapters hijack the RNA helicase UPF1 in a CRM1/XPO1 dependent manner and reveal proviral roles of UPF1"

### Supplementary data

Supplementary Figure 1

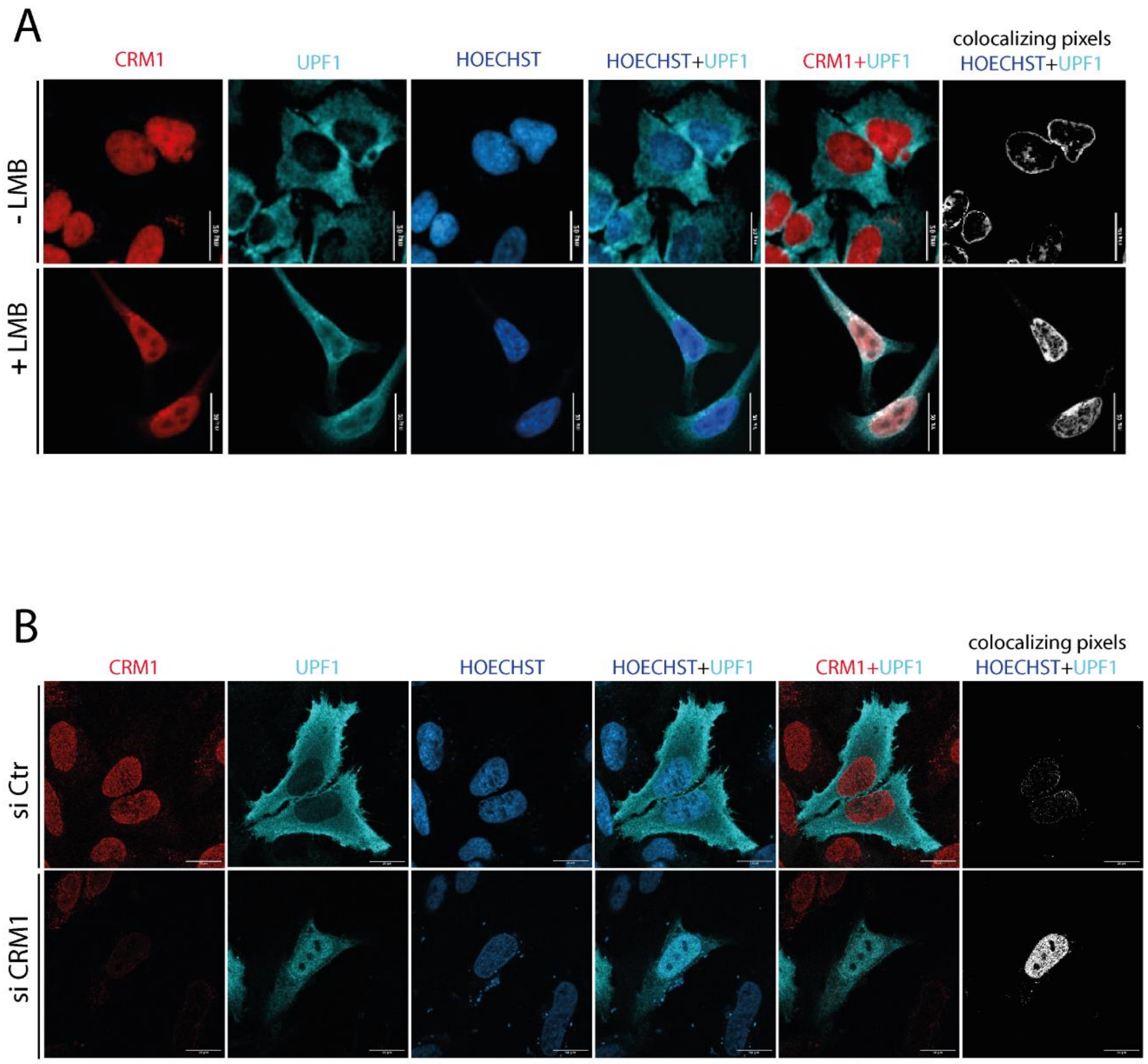

**Supplementary Fig1 (relative to Fig1)**:

A) 6h before fixation, HeLa cells were treated with either 50nM leptomycin B (LMB, an inhibitor of CRM1 export) or EtOH as a control. Cells were further stained with antibodies against CRM1 and UPF1. Nucleus were stained with Hoechst. Merge with UPF1 and Hoechst or UPF1 and CRM1 were performed as indicated. The pixels excited with anti UPF1 and Hoechst staining simultaneously are shown as “Colocalizing pixels”. Objective x63. Scale bar : 30µm. This experiment shows a nuclear retention of endogenous UPF1 after LMB treatment.

B) HeLa cells were transfected with 10nM of siRNA control or against CRM1 for 72h. Cells were analysed similarly as in A. SiCRM1 induced a strong reduction of CRM1 levels. Endogenous UPF1 localization is drastically modified after the treatment with siCRM1 displaying nuclear retention. Objective x63. Scale bar : 20µm

Altogether, these results confirm data from the literature, indicating that UPF1 export is mediated by CRM1.

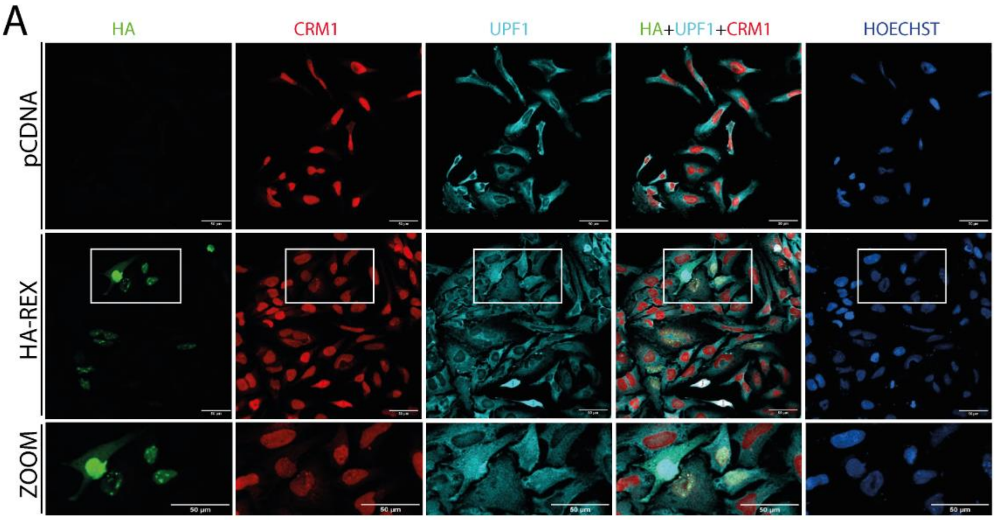

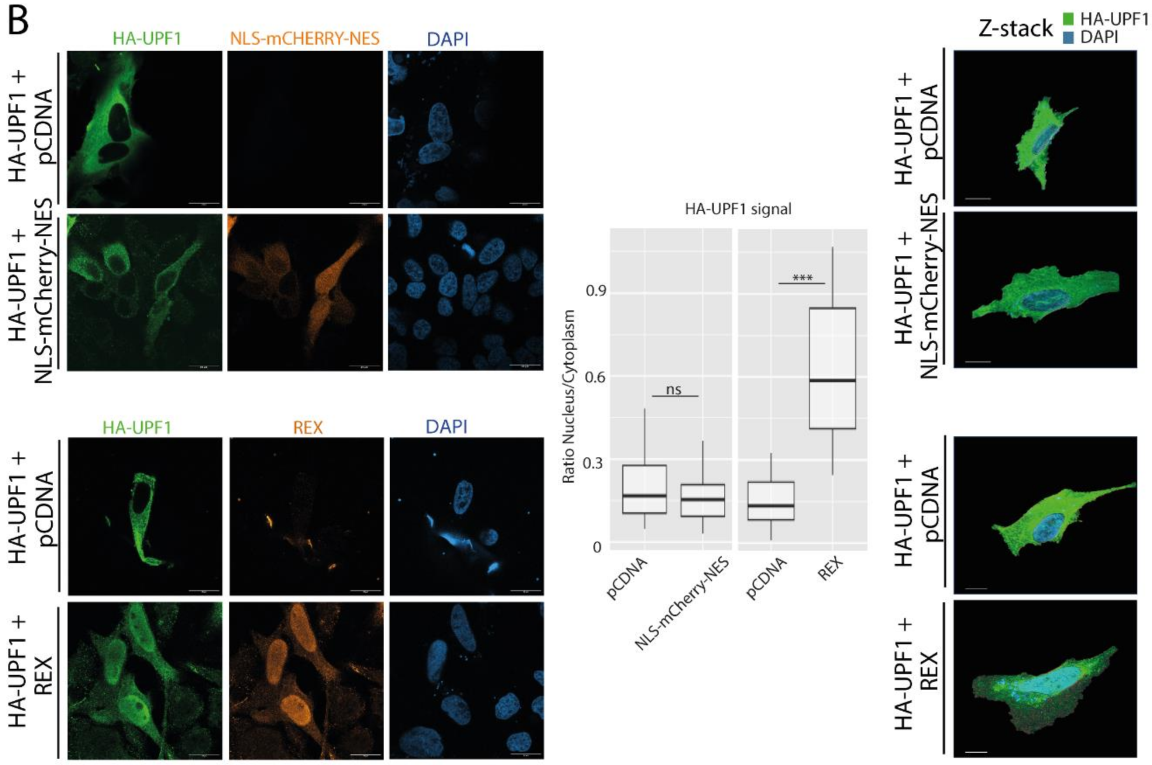

**Supplementary Fig2 (relative to Fig1):**

A) Wide field view of the confocal microscopy experiment showing the nuclear accumulation of UPF1 induced by Rex overexpression (related to **Fig 1C)**. Magnification of the framed zone is shown in the lower panel (ZOOM). Objective x20. Scale bar: 50µm. B) (left panel) Confocal microscopy of HeLa cells transfected with the indicated combinations of HA-UPF1, NLS-mCherry-NES and Rex coding plasmids. HA-UPF1 and mCherry were revealed. Objective x63. Scale bar: 20µm. (middle panel) Quantitative analysis of the microscopy experiments: mCherry signal was quantified in the cytoplasm and in the nucleus and expressed as a ratio. Wilcoxon, pvalue ns> 0.05; ***<0.005. (right panel) Z-stack analysis of UPF1 staining. These experiments show that the modification of UPF1 localisation profile is specific of Rex and cannot be mimic by any overexpressed NES-containing protein. As a consequence, the nuclear retention of UPF1 induced by Rex is not based on a competition to bind CRM1.

Supplementary Figure 3

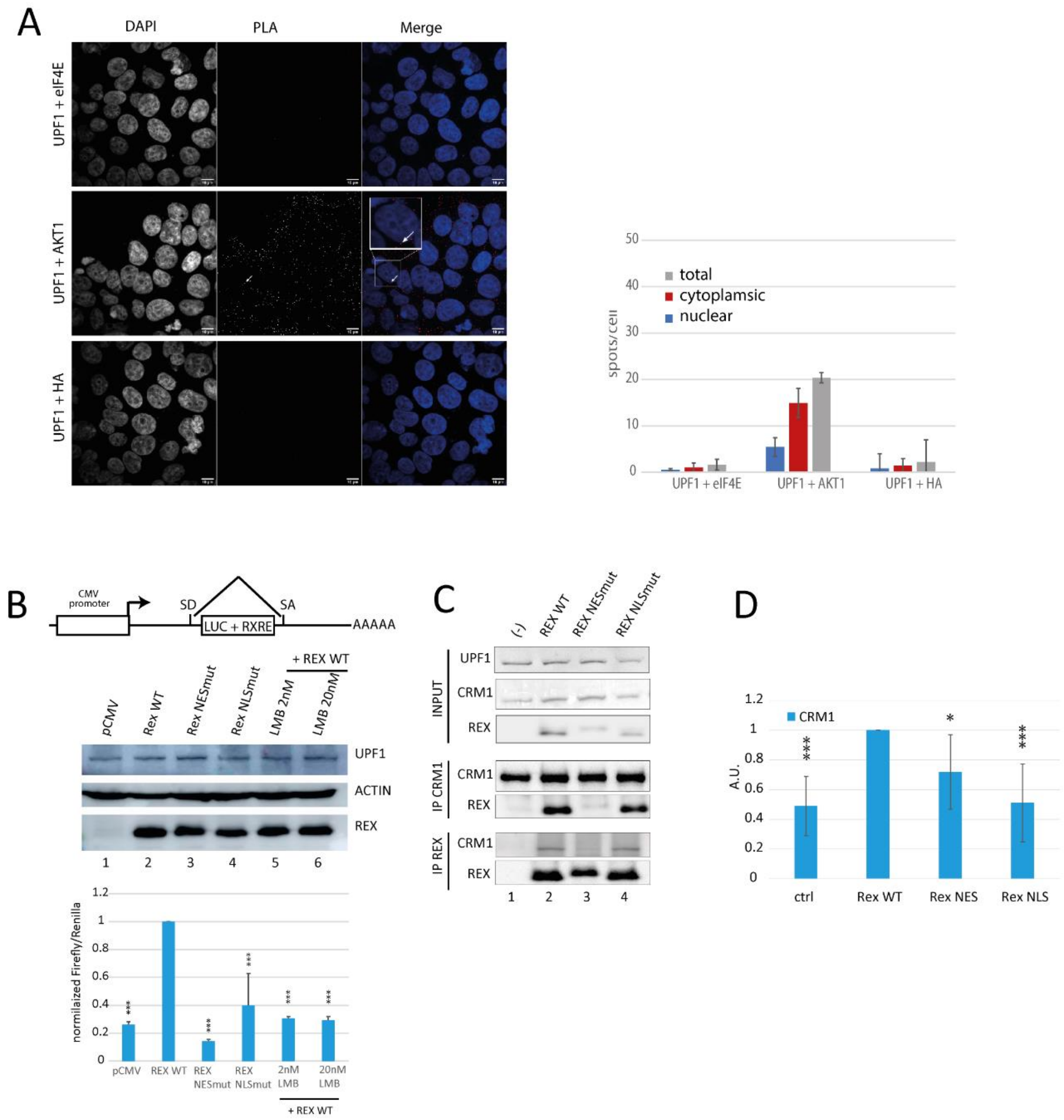

**Supplementary Fig3 (relative to Fig 2)**:

A) PLA control experiments from Fig 2A, showing UPF1 signal specificity. PLA were performed using anti UPF1 combined with anti eIF4E or anti HA. No spots are generated, as expected of negative controls. Anti UPF1 combined with anti AKT1 generated spots as shown previously(1). White arrow pinpointes one example of dot. B) To confirm and further characterize the role of Rex in favouring UPF1-CRM1 interaction, we carried out experiments with cells expressing Rex mutants. Rex dependent RNA export assay was performed as previously described(2). The Rex dependent RNA reporter is composed of the luciferase gene followed by the RxRE motif under the control of a pCMV promoter. The luciferase/RxRE cassette is encompassed with splicing donor and acceptor sites that provoke its splicing preventing luciferase expression in the absence of Rex (upper panel). When Rex is expressed, this latter binds the RxRE, prevents the splicing of the Luciferase/RxRE cassette and drives its export in a CRM1 dependent manner (like with vRNA), allowing luciferase expression. The luciferase expression is quantified with the "luciferase dual GLO luciferase assay" (Promega). A Renilla coding plasmid is co-transfected with the luc-RxRE plasmid for normalization, following the manufacturer protocol. In addition, we transfected Rex WT (positive control, lane2), Rex NES mutant (lane3), Rex NLS mutant (lane 4) or an empty vector (negative control, lane1). As controls we also treated cells that expressed Rex with LMB at the indicated concentrations for 12h (lanes 5-6). Protein level expression was controlled by western blot (middle panel). Luciferase quantification were displayed as a percentage of the Rex WT condition (lower panel, lane2). As expected, in the absence of Rex or when cells were treated with LMB, luciferase expression was strongly reduced (lower panel, compare lane1, 5, 6 and 2). The expression of Rex NES mutant or Rex NLS mutant also impaired luciferase expression, validating the export defect of both mutants (lower panel, compare lane 3-4 and 2). C) coIP experiment in 293T cells transfected with WT and mutant forms of Rex coding plasmids. Immunoprecipitations of either CRM1 or Rex show that Rex NES mutant is not able to interact with CRM1 (lane 3) compared with Rex WT (lane 2), while Rex NLS mutant is still able (lane 4). D) Quantification of the western blots from **Fig 2E**. Bars represent the medium of 4 independent experiments. Co-immunoprecipitated CRM1 was quantified using ImageLab (Biorad) and further normalized by the levels of immunoprecipitated UPF1(IP UPF1) and the levels of CRM1 in the total extracts (input). Rex with a mutation in the Nuclear Export Signal (NES) no longer interacts with CRM1 and is functionally not able to export RNA containing a RxRE motif compared to Rex WT; Rex with a mutation in the Nuclear Localisation Signal (NLS) still interacts with CRM1 but is also greatly impaired in RNA export (**supplementary Fig3B, C**). Noteworthy, the NLS mutation affects the UPF1/Rex association, as shown in UPF1 coIP experiments (**Fig 2E**).

Supplementary Figure 4

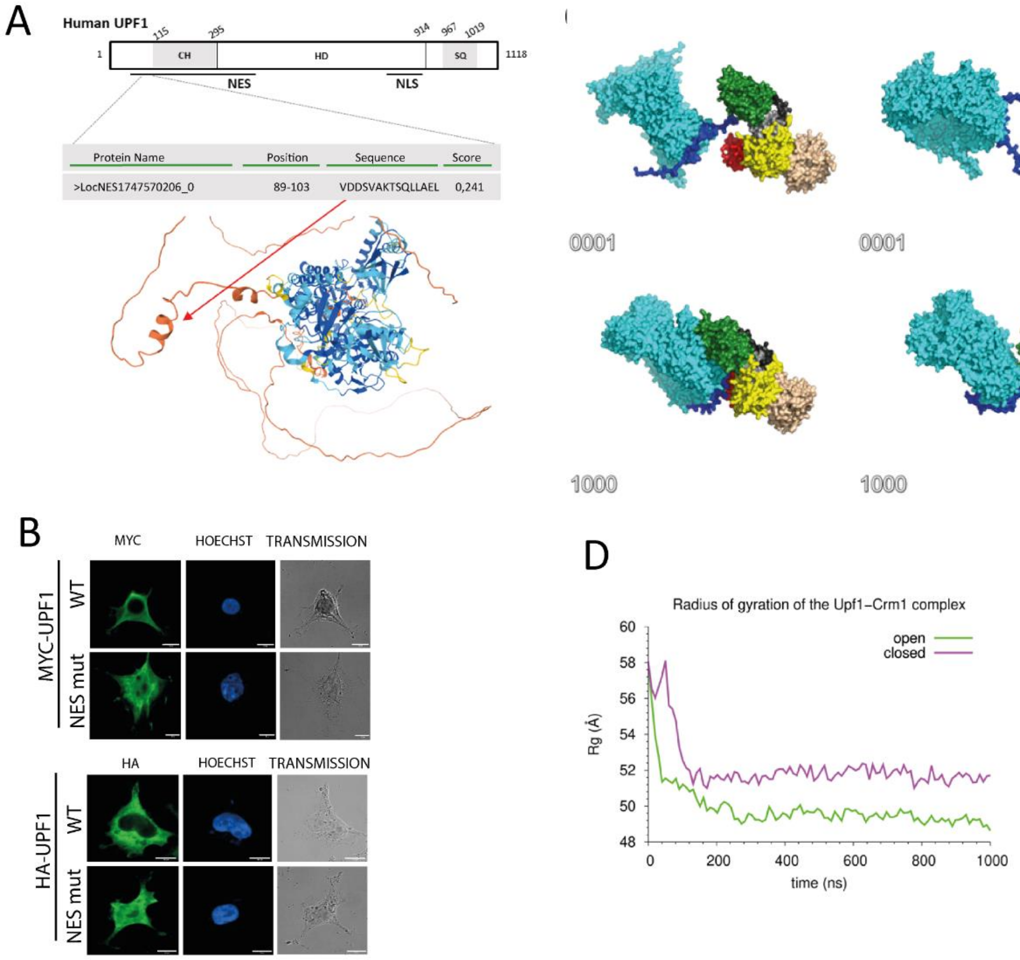

**Supplementary Fig4 (relative to Fig 3)**:

A) Schematic representation of UPF1 protein with the NES and NLS delineation from the literature (in black), the precise NES sequence identified using LocNES software and its positioning using a alfafold2 modelling of UPF1 full-length. B) To confirm the function of this putative NES sequence, a plasmid coding a UPF1 NES mutant (VDDSVAKTSQLLAEL -> VDDSVAKTSQG) was engineered: to the contrary of the UPF1 WT, UPF1 NES mutant displays a clear nuclear accumulation, demonstrating an export defect. The mutation was incorporated in two different backbones: HA-UPF1 and Myc-UPF1. Myc-UPF1 was modified with silent mutations to make UPF1 resistant to siUPF1. C) Coarse-grained molecular modelling with Martini force field of CRM1-UPF1 complex starting from UPF1 which CH domain open (PDB 2WJV) of UPF1 with CH domain closed (PDB 2XZL). Initial (0001) and final (1000ns) states of simulations of UPF1-CRM1 complexes. CRM1 is colored in cyan. UPF1 NES (blue), CH (green), 1B (red), 1C (orange), RecA1 (yellow), RecA2 (wheat). D) Monitoring of the Radius of gyration of the UPF1-CRM1 complex during the 1000ns of simulation.

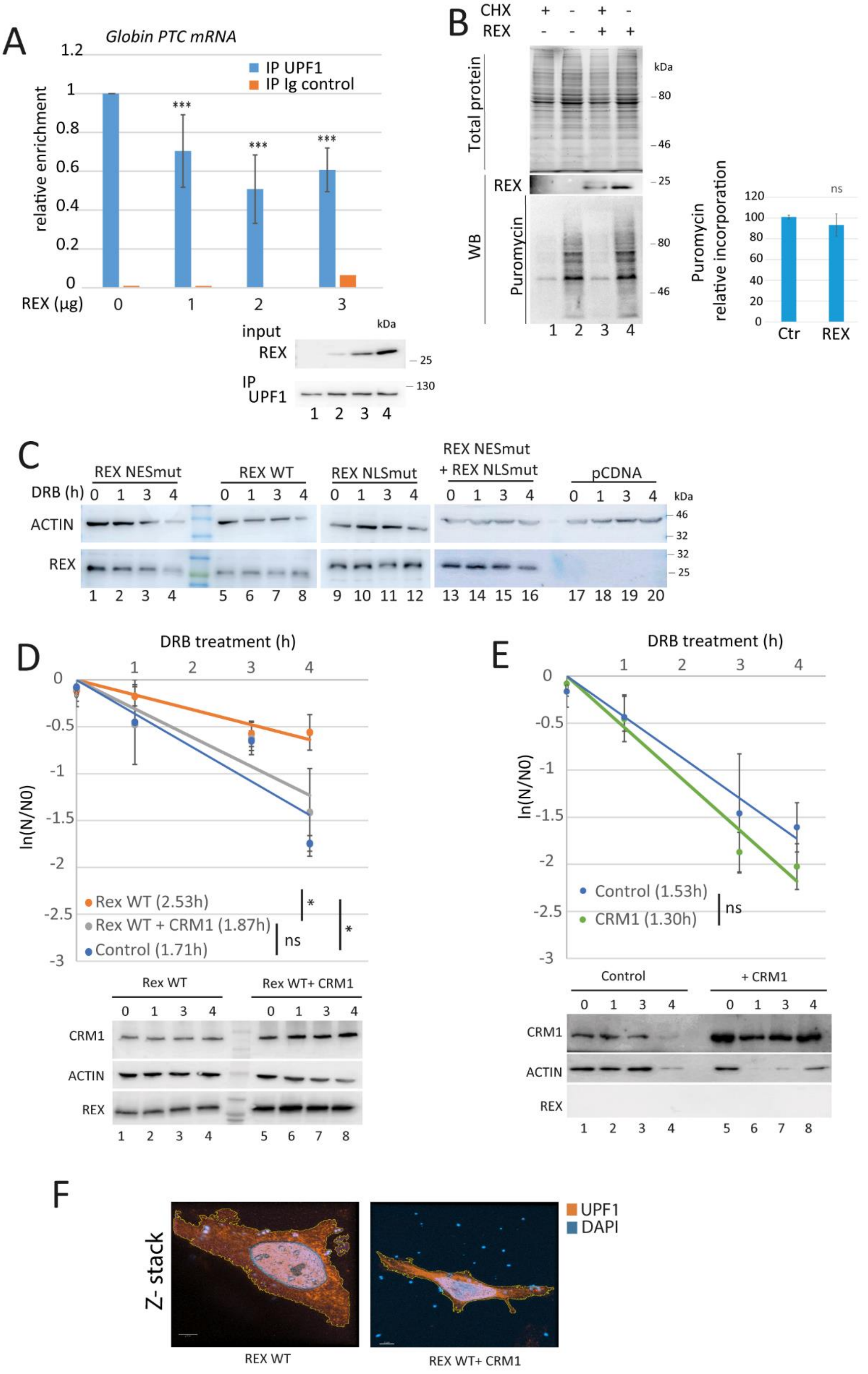
A
Globin PTC mRNA
relative enrichment
IP UPF1
IP Ig control
REX (µg)
input
REX
IP UPF1
kDa
B
CHX
REX
Total protein
WB
Puromycin
Puromycin relative incorporation
Ctr
REX
ns
C
REX NESmut
REX WT
REX NLSmut
REX NESmut + REX NLSmut
pCDNA
DRB (h)
ACTIN
REX
D
DRB treatment (h)
ln(N/N0)
Rex WT (2.53h)
Rex WT + CRM1 (1.87h)
Control (1.71h)
Rex WT
Rex WT+ CRM1
CRM1
ACTIN
REX
E
DRB treatment (h)
ln(N/N0)
Control (1.53h)
CRM1 (1.30h)
Control
+ CRM1
CRM1
ACTIN
REX
F
Z- stack
UPF1
DAPI
REX WT
REX WT+ CRM1

**Supplementary Fig5 (relative to Fig 3)**:

A) RNA immunoprecipitation experiment (RIP) using a rabbit polyclonal antibody targeting UPF1 (blue) or from a pre-immune serum (Ig control, orange) were performed with HeLa cells transfected with a Globin PTC coding plasmid and increasing amounts of Rex coding plasmid (0, 1, 2, 3µg). Immunoprecipitated RNA (Globin PTC) were quantified by RTqPCR. For each condition, the relative enrichment of Globin PTC RNA associated to UPF1 compared to conditions without Rex was displayed in the graph. On the right panel, western blot controlling the levels of the expressed Rex and the immunoprecipitated UPF1. A dose response effect was observed up to 2µg of transfected Rex. Above, the concentration had an inhibiting effect. We choose to transfect 1.5µg in the following experiments. Ttest pvalue: ***<0.005. B) Sunset assay with HeLa cells expressing or not Rex (lane 2 and 4). The level of total protein was acquired before transfer with stainless method, Rex expression and puromycin incorporation were evaluated by western blot. Puromycin quantification was indicated on the right panel. As a negative control, cells were pre-treated with cycloheximid to inhibit translation (lane 1 and 3). Statistical analysis: ttest ns: $p>0.05$. C) Western blot controls from **Fig3D**. D) Half-life evaluation of the Glob PTC mRNA in HeLa cells transfected with a control plasmid, Rex or Rex complemented with CRM1. mRNA half-lifes ($t1/2 = \ln(2)/\lambda$ with $\lambda$ the time constant of the decay curves) are indicated in front of their respective conditions. ttest pvalue: ns>0.05; *< 0.05. Proteins expression were controlled by western blot. E) Same as D with Hela cells overexpressing CRM1 or not, in the absence of Rex. F) Z stack analysis from the confocal microscopy experiments presented in **Fig3F**. The conditions of UPF1 nuclear retention in presence of Rex and the reversion of the phenotype with CRM1 overexpression are presented.

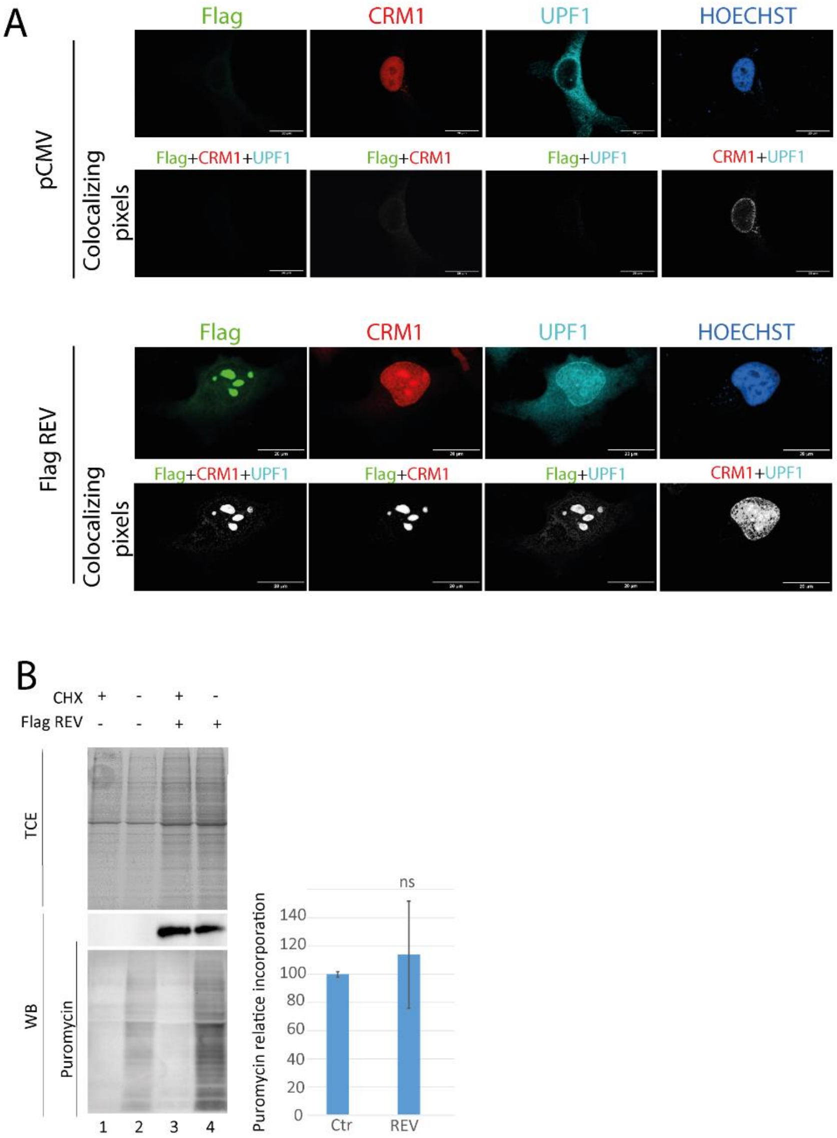

**Supplementary Fig6 (relative to Fig 4)**:

A) Confocal microscopy experiments were performed in HeLa cells transfected with the Flag-Rev plasmid. Rev was revealed with an anti-Flag tag antibody and an alexa 488 secondary anti-mouse. CRM1 antibody was targeted with an alexa 647 anti-rabbit and the UPF1 antibody with a CFL 555 anti-goat. The colocalizing pixels view is obtained as mentioned in the methods section and only show the pixels activated in common at the indicated conditions. Images were acquired with a x63 objective. Scale bar: 20µm. On this representative close up image, we clearly observed the nuclear localisation of UPF1 and its colocalization with Rev and CRM1. B) Sunset assay with HeLa cells expressing or not Rev (lane 2 and 4). The level of total protein was acquired before transfer with a stainless method, Rex expression and puromycin incorporation were evaluated by western blot. Puromycin quantification was indicated on the right panel. As a negative control, cells were pre-treated with cycloheximid to inhibit translation (lane 1 and 3). Statistical analysis: Ttest ns: $p>0.05$.

Supplementary Figure 7

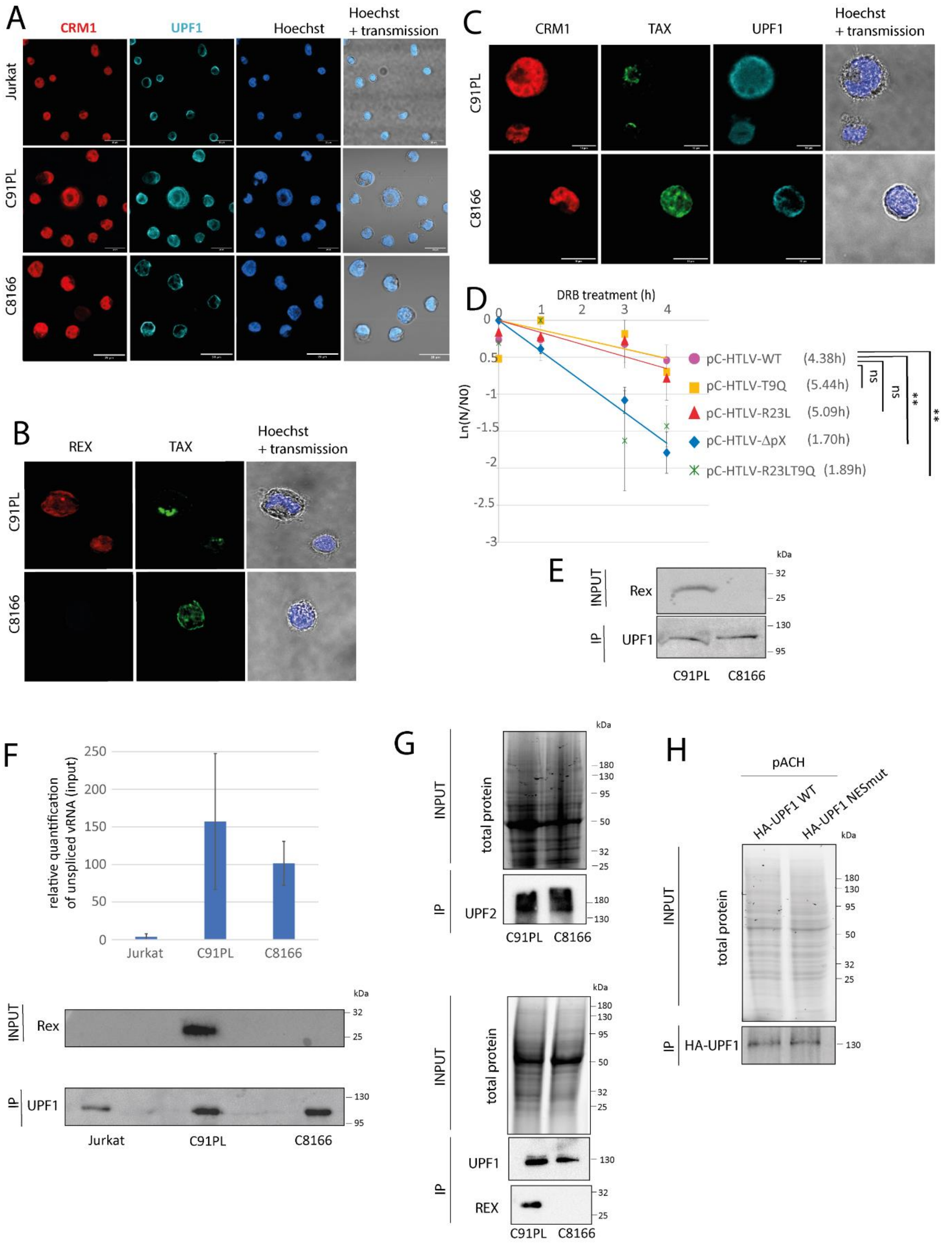

**Supplementary Fig7 (relative to Fig 5-6)**:

A) Wide field views of Jurkat, C91PL and C8166 lymphocytes stained for CRM1 and UPF1 and observed by confocal microscopy. Images were acquired with a x63 objective. Scale bar: 20µm. B) Close up view of C91PL and C8166 lymphocytes stained for Rex and Tax and

observed by confocal microscopy. Images were acquired with a x63 objective. Scale bar: 10µm
C) same as B with CRM1, Tax and UPF1 staining.
D) Half-life evaluation of the Glob PTC mRNA in HeLa cells transfected with the indicated forms of HTLV-1 molecular clones:
-pC-HTLV-WT
-pC-HTLV-T9Q (extinction of Tax expression due to a point mutation inducing nonsense codon at position 9 of Tax but maintaining Rex expression)
-pC-HTLV-R23L (extinction of Rex expression due to a point mutation inducing a nonsense mutation at position 90 of Rex but maintaining Tax expression)
-pC-HTLV-R23LT9Q (extinction of Tax and Rex expression due to the combination of the 2-point mutations)
-pC-HTLV-ΔpX (extinction of Tax and Rex expression due to the deletion of ~920nt across the pX region).

mRNA half-lifes were measured as described in Fig3 and 4, ($t1/2 = \ln(2)/\lambda$ with $\lambda$ the time constant of the decay curves) and reported in front of their respective conditions. Statistical analysis: Ttest ** $p<0.01$; ns : $p>0.05$.

The pC-HTLV molecular clone has the 5'LTR sequence replaced by a CMV promoter. Consequently, the virus transcription is not dependant on Tax expression. pC-HTLV-WT is able to inhibit NMD while pC-HTLV-ΔpX couldn't, as shown previously (3). To evaluate the specific impact of Rex on NMD without Tax interference and in a context a virus expression, we used the pC-HTLV-T9Q mutant where Tax ORF was mutated to prevent its expression without altering Rex and viral transcription. In addition, we engineered the pC-HTLV-R23LT9Q that lacks Tax and Rex expression. Regarding Rex specific effect on NMD, pC-HTLV-T9Q and pC-HTLV-R23LT9Q could be considered as mimicry of C91PL and C8166 cell lines, respectively.
As expected, Glob PTC half-life measurement showed that pC-HTLV-T9Q inhibits NMD while pC-HTLV-R23LT9Q couldn't. This validates the capacity of Rex to inhibit NMD, independently of Tax, in a context of virus expression.

E) Western Blot control for RIP experiment **Fig5D**. F) Bar plot representation of the relative quantification of the unspliced viral RNA in Jurkat, C8166 and C91PL total extracts used to carry out RIP in **Fig6A**. Western blot controls for the RIP experiment **Fig 6A**. G) Western Blot control for RIP experiment **Fig6B**. Total protein input is visualized with stainfree gels.
H) Western Blot control for RIP experiment **Fig 6F**. Total protein input is visualized with stainfree gels

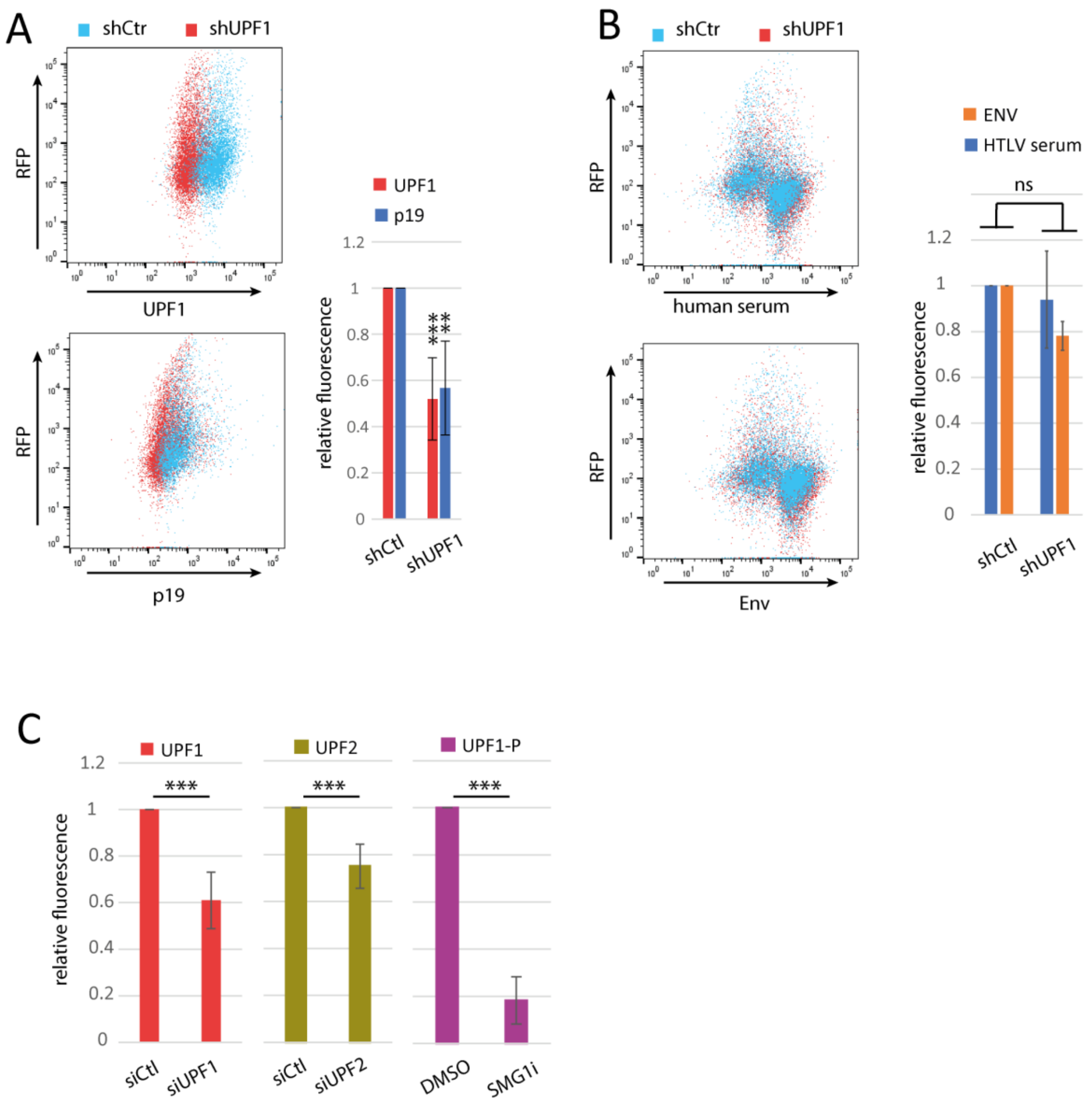

**Supplementary Fig8 (relative to Fig 7)**:

To evaluate the impact of UPF1 on the production of structural proteins, cellular samples from **Fig 7B** were also monitored by FACS. A) Cells were washed, fixed and permeabilized before immunostaining against UPF1 or MA/p19. From RFP positive cells (expressing shRNA constructions), the mean fluorescence of UPF1 or MA/p19 labelling was quantified and is shown in the bar plot. We found that UPF1 extinction was associated with a significant decreased of MA/p19 levels, as described in Fig7B(i). B) In order to evaluate the levels of the Env protein at the surface of the producing cells (representative of the amount of membrane-bound viral particles), 293T cells were stained with an anti HTLV-1 serum (gift from A.Gessain) or an anti Env antibody, without permeabilization and before fixation. Under these conditions, we found a small non-significant decrease of the signal in the absence of UPF1, confirming the western blot observations in Fig7B(ii). C) Control of the siRNA treatments and the inhibition of UPF1 phosphorylation by SMG1i (relative to Fig7G).

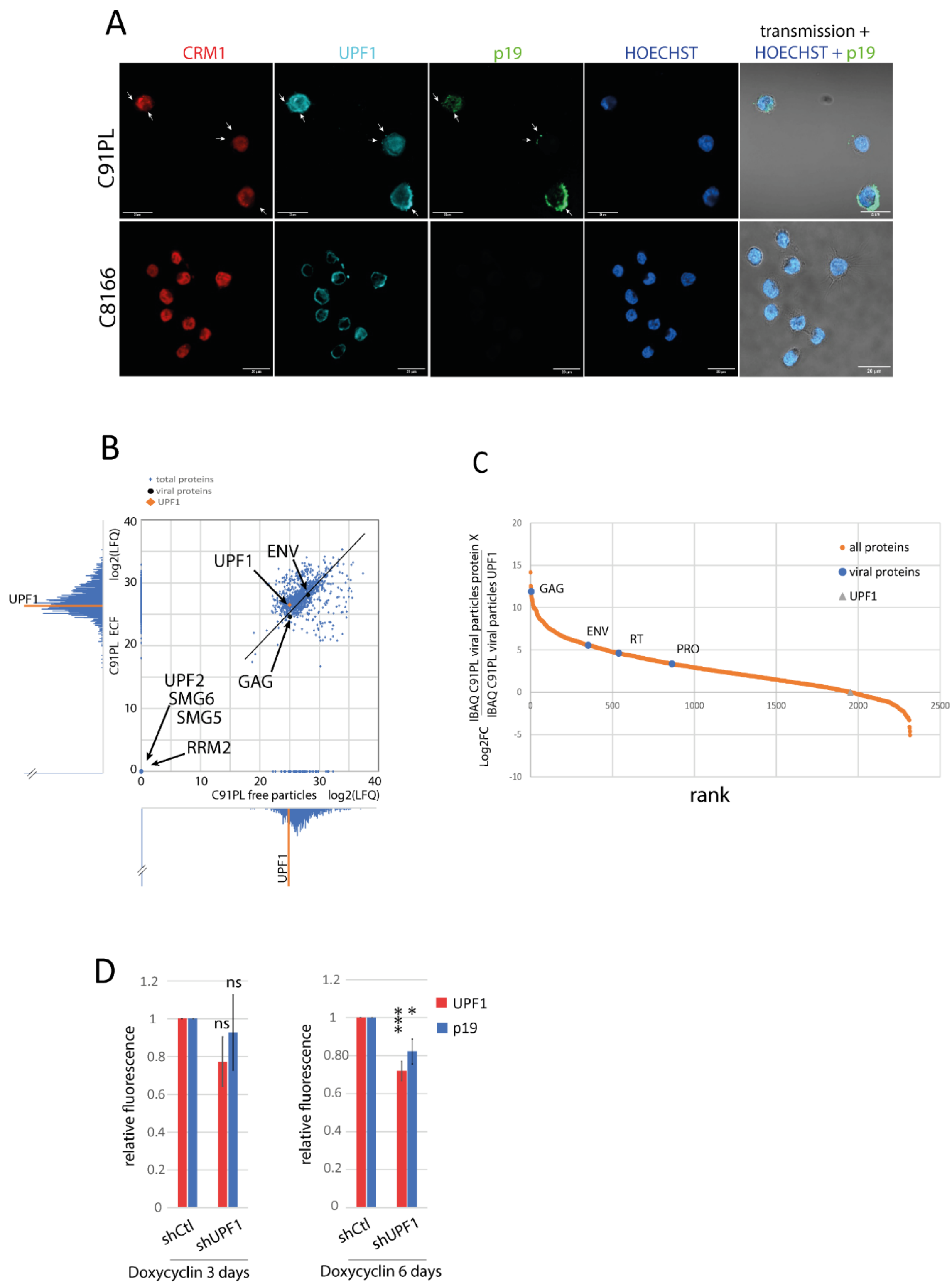

**Supplementary Fig9 (relative to Fig 8)**:

A) Wide field views of C91PL and C8166 lymphocytes stained for CRM1, UPF1 and P19 observed by confocal microscopy (related to **Fig 6A).** B) Distribution of the proteins quantified by mass spectrometry in the "free particles" fraction (x axis) and in the ECF fraction (y axis) of C91PL cultures. Each coordinate corresponds to the mean of 3 independent experiments. UPF1 position in the scatter plot is represented in orange. Viral proteins GAG and ENV were found in the C91PL free particles only, validating the enrichment of virions in this fraction. C) We evaluate the stochïometry of UPF1 compared to the other components of the viral particles from C91PL cultures. UPF1 is identified as a relatively low concentrated protein in the HTLV-1 viral particles. D) C91PL were infected with lentivirus expressing shRNA against UPF1 or control, as indicated in Fig.7H. The levels of MA/p19 and UPF1 of C91PL after 3 or 6 days of shRNA induction by doxycyclin treatment were assessed by FACS. The mean fluorescence is represented in the bar plot. This shows that a significant decrease of UPF1 was achieved after 6 days of shRNA induction. This decrease was associated with a significant drop in MA/p19 levels.

**Supplementary methods**

**Antibodies**. Anti-UPF1 rabbit polyclonal (RENT1, A301-902A, Bethyl), Anti-CRM1 rabbit polyclonal (A300-469A, Bethyl), Anti-UPF2 rabbit polyclonal (A303-029A, Bethyl), Anti-HA mouse monoclonal (clone HA-7, SIGMA), Anti-FLAG mouse monoclonal (FlagM2, Sigma), Anti-HTLV-1 p19 mouse monoclonal (TP-7, Abcam), Anti-Tax mouse monoclonal, Anti-Rex rabbit polyclonal(4) and Anti-Ribonucleoside-diphosphate reductase subunit M2 (RRM2) rabbit polyclonal(5). For immunofluorescence specifically: primary Goat UPF1 (A300-036A, Bethyl), and secondary antibodies were goat anti-mouse Alexa Fluor 488 (Thermofischer), mouse anti-Goat CFL 555 (Santa Cruz), Donkey anti-Rabbit Alexa Fluor 647 (Abcam).

**Primers**

| name | Forward (5'->3') | Reverse (5'->3') |
|---|---|---|
| **Mutagenesis** | | |
| Rex NLSmut | cgatcccaaagaGATTTAccaccaacacc | ggtgttggtggTAAATCtctttgggatcg |
| Rex NESmut1 | tcagctctacagttccGGAtcccctccttcc | ggaaggaggggaTCCggaactgtagagct ga |
| Rex NESmut2 | ctacagttccGGAtcccctccttccccac | gaaggaggggaTCCggaactgtagagctg |
| pCMV HTLV Rex 23L | ggacgcgttatcagctcagctctacagttccTAAtc cctcg | atttgtcttcagggggacac |
| **RTqPCR** | | |
| globin | TTGGGGATCTGTCCACTCC | CACACCAGCCACCACTTTC |
| renilla | CTAACCTCGCCCTTCTCCTT | TCGTCCATGCTGAGAGTGTC |
| viral unspliced RNA (vRNA) | GGCCCGAGGACACACTAATA | CAGCGGGGAGGTCTAATAGG |
| GADD45α | ACGAGGACGACGACAGAGAT | GCAGGATCCTTCCATTGAGA |
| GAPDH | GAGTCAACGGATTTGGTCGT | TTGATTTTGGAGGGATCTCG |
| SMG5 | ACAGAATGGGATGCCAGGAA | TCAACAC TCCAAAAGCCAGC |

**Cell fractionation**. ~$2\times10^6$ HeLa cells were transfected as described in the figures, harvested and resuspended in lysis buffer (340 mM Sucrose, 10% Glycerol, 10mM HEPES pH 7.5, KCl, 1.5mM MgCl2, 0.1% Triton, 1mM DTT, 1mM Na3VO4, protease inhibitor (Roche) and RNAsin (Promega)). The cytoplasmic fraction was collected after 1300g centrifugation and conserved for further RNA extraction with RNAzol RT reagent (MRC) and subjected to RT-qPCR or for Western blotting. The isolated nucleus were washed two times with lysis buffer and further pelleted. Nucleus were resuspended in nucleus lysis buffer (3mM EDTA, 0.2mM EGTA, 0.1% Triton, 1mM DTT, 1mM Na3VO4, protease inhibitor (Roche) and RNAsin (Promega))) and kept under rotation during 30min at 4°C. Nuclear fraction was collected after 12 000g centrifugation. Total RNA was extracted with RNAzol RT and subjected to qRT-PCR. The values represented in the graphs correspond to the mean of at least three biological replicates, and the error bars correspond to the SD. P values were calculated by performing a Student's t-test (unpaired, two-tailed) ns: $P > 0.05$; $*P < 0.05$; $**P < 0.01$

**FACS**

FACS analysis were performed on 293T cells transduced with the Dharmacon SMARTvector inducible shRNA (either control or targeting UPF1). Cells were placed under puromycin selection and shRNA were induced with doxycycline (0.5µg/ml) as indicated. 48h after the transfection of a HTLV-1 molecular clone, cells were washed and fixed with paraformaldehyde 4% for a further FACS analysis(6). Cells were permeabilized, divided in two batches, each one being incubated with primary antibodies either against UPF1 (rabbit polyclonal) or against the viral matrix (p19) (mouse monoclonal). Alexa 488 secondary antibodies were then used. 5000-10000 cells were analysed by flow cytometry analysis, which was carried out with a MACSQuant VYB apparatus equipped with 488- and 561-nm emission lasers (Miltenyi Biotec, Bergisch Gladbach, Germany): cells were first selected based on RFP expression (RFP is constitutively co-expressed with shRNA after doxycycline induction in the Dharmacon SMARTvector inducible plasmids). Then cells were selected based on UPF1 or p19 labelling and the mean fluorescence was quantified and represented in bar plot. Background fluorescence was evaluated by the same analysis on non-transduced and non-transfected 293T. For Env quantification, the same protocol was followed except that cells were not permeabilized and that the primary antibodies' incubation was done prior fixation to prevent intracellular staining.

**Metabolic assays (SUnSET®).** The SUnSET® assay was used to monitor de novo protein synthesis as described previously (7). ~$2\times10^6$ of HeLa cells were transfected as described in the figures. Briefly, 10min prior harvesting the cells, puromycin was added to culture medium at 1µg/ml. As a control, cycloheximide was added at 10µg/ml, 15min before puromycin addition, resulting in complete blockade of protein synthesis. Cell extracts were then processed for Western blotting using anti-puromycin 12D10 antibody (Millipore) at $1/5000^{eme}$. Puromycin relative incorporation was calculated based on quantification of puromycin signals normalized by total protein. Total protein levels were visualized with stain free gels. Western blot signals and total protein levels were acquired on a Biorad Chemidoc imaging system before quantification with Image Lab software. The values represented in the graphs correspond to the mean of at least three biological replicates, and the error bars correspond to the SD. P values were calculated by performing a Student's t-test (unpaired, two-tailed) ns: $P > 0.05$; $*P < 0.05$; $**P < 0.01$.

**Supplementary references:**

1. Palma,M., Leroy,C., Salomé-Desnoulez,S., Werkmeister,E., Kong,R., Mongy,M., Le Hir,H. and Lejeune,F. (2021) A role for AKT1 in nonsense-mediated mRNA decay. *Nucleic Acids Res.*, **49**, 11022–11037.
2. Kress,E., Baydoun,H.H., Bex,F., Gazzolo,L. and Duc Dodon,M. (2005) Critical role of hnRNP A1 in HTLV-1 replication in human transformed T lymphocytes. *Retrovirology*, **2**, 8.
3. Fiorini,F., Robin,J.-P., Kanaan,J., Borowiak,M., Croquette,V., Le Hir,H., Jalinot,P. and Mocquet,V. (2018) HTLV-1 Tax plugs and freezes UPF1 helicase leading to nonsense-mediated mRNA decay inhibition. *Nat. Commun.*, **9**.
4. Dodon,M.D., Hamaia,S., Martin,J. and Gazzolo,L. (2002) Heterogeneous Nuclear Ribonucleoprotein A1 Interferes with the Binding of the Human T Cell Leukemia Virus Type 1 Rex Regulatory Protein to Its Response Element. *J. Biol. Chem.*, **277**, 18744–18752.
5. Kuo,M. and Kinsella,T. (1997) Overexpression of a hexa-histidine and T7 peptide-tagged human ribonucleotide reductase small subunit, R2 in Escherichia coli and the generation of human R2 antibodies. *Int. J. Oncol.*, **10**, 515–520.

6. Roisin,A., Buchsbaum,S., Mocquet,V. and Jalinot,P. (2021) The fluorescent protein stability assay: an efficient method for monitoring intracellular protein stability. *BioTechniques*, **70**, 336–344.
7. Schmidt,E.K., Clavarino,G., Ceppi,M. and Pierre,P. (2009) SUnSET, a nonradioactive method to monitor protein synthesis. *Nat. Methods*, **6**, 275–277.
